# Toehold-VISTA: A machine learning approach to decipher programmable RNA sensor-target interactions

**DOI:** 10.1101/2025.08.12.669990

**Authors:** James M. Robson, Alexander A. Green

## Abstract

RNA-based biosensors have emerged as essential tools in synthetic biology and diagnostics, enabling precise and programmable responses to diverse RNA inputs. However, the time to design, produce, and screen high-performance RNA sensors remains a critical challenge. The fundamental rules governing RNA-RNA interactions—specifically the structure-function relationships that determine sensor performance—remain poorly understood. Here, we present a method enabling versatile in-silico RNA-targeting analysis (VISTA), a machine learning-guided framework for the rapid design of RNA sensors. VISTA integrates biophysical modeling of both sensor and target RNAs with a partial least squares discriminant analysis (PLS-DA) machine learning framework. Using high-throughput experimental measurements with sequence-structure feature extraction to train predictive models, we capture the key determinants of RNA sensor performance. We find that by using toehold switches as a model RNA sensor, Toehold-VISTA successfully designs RNA sensors with improved function against SARS-CoV-2 RNA. These findings establish a broadly applicable, target-aware design strategy for accelerating RNA sensor engineering across biotechnology and diagnostic applications.

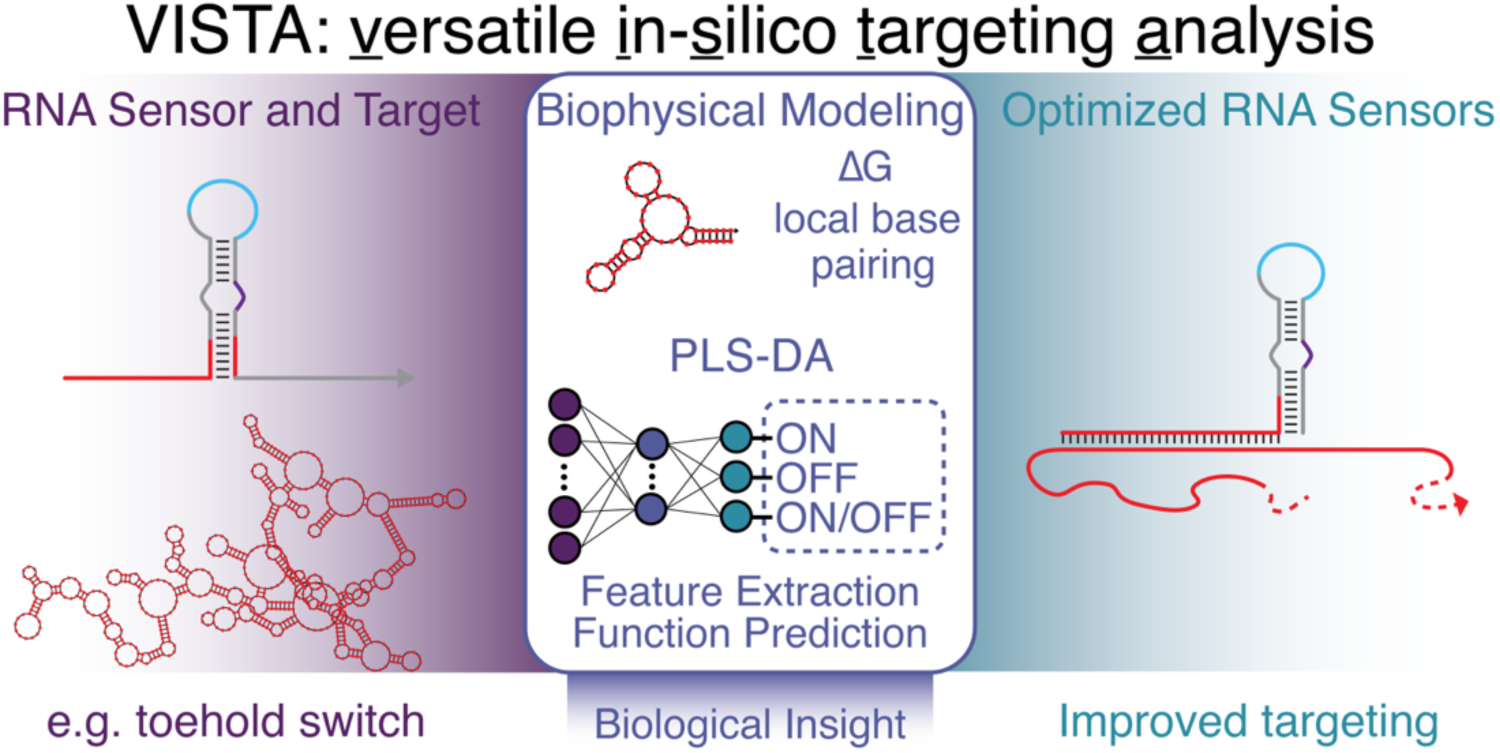

## INTRODUCTION

RNA-based regulators are transformative tools in synthetic biology, diagnostics, and biotechnology, enabling programmable detection of nucleic acid inputs and precise control of gene expression (1–7). These systems capitalize on the predictable base-pairing rules of RNA to sense specific RNA or DNA sequences, converting this molecular recognition into a functional output (8–18). RNA sensors offer advantages in terms of modularity, rapid programmability, and compatibility with both cell-free and in vivo platforms, making them highly attractive for applications from environmental biosensing to clinical diagnostics (19–26). However, the performance of these sensors is heavily influenced by the often-overlooked interplay between the sensor and its target RNA, particularly in trans-acting systems where binding kinetics, secondary structure, and accessibility dictate function (2, 5, 6, 9, 19).

Despite the rapid development of design tools and predictive models for RNA-based sensors (27, 28), most frameworks have focused primarily on features intrinsic to the sensor RNA, such as folding stability, ribosome accessibility, or thermodynamic properties of hairpin formation. Prior studies have examined aspects of target RNA folding, but few have systematically or comprehensively evaluated how these structural features affect sensor performance in empirical settings. Enhancing RNA sensor functionality requires a deeper understanding of target RNAs— including viral genomes, endogenous transcripts, and synthetic constructs—and the complex secondary structures they form, which can mask sensor-binding sites and hinder detection. Work in CRISPR-Cas guide RNA and microRNA target design has underscored the importance of target site accessibility for effective binding and function, yet this critical consideration has not been explored in the context of other RNA sensors (29–33).

Most current models for RNA sensor design, including recent deep learning-based predictors (34–36), fail to incorporate features derived from the structural context of the RNA target, including local base-pairing probabilities, minimum free energy, and codon usage bias that may impact co-transcriptional folding (37, 38). When sensing full-length RNA transcripts, this limitation is particularly pronounced as distal structural elements can alter the folding landscape and prevent effective hybridization at the intended target site. The need for target-aware design principles is thus critical for improving the robustness and generalizability of RNA sensor design, especially to reduce the labor and time required for the design-build-test cycle in RNA synthetic biology.

To address this gap, we systematically investigated the role of trans-RNA structure-function relationships by pairing a comprehensive library of toehold switch riboregulators with tiled binding sites across a structured RNA transcript in *E. coli*. By combining biophysical modeling of both the toehold switch and its trans-RNA target, we quantified key features—including local and global structural accessibility—and their influence on sensor performance. We then applied a partial least squares discriminant analysis (PLS-DA) machine learning framework to identify the most informative features for predicting sensor activity across multiple functional metrics. The resulting platform, versatile in-silico RNA-targeting analysis, or VISTA, incorporates both sensor and target design parameters into a unified RNA sensor design pipeline. Employing VISTA for the design of toehold switches (Toehold-VISTA), we dramatically improve prediction accuracy over existing toehold switch design tools. We establish a generalizable strategy for RNA sensor development that highlights the importance of trans-RNA interactions and structural context as key design principles for engineering nucleic acid sensors with enhanced robustness and performance.

## MATERIAL AND METHODS

### Design of toehold switches

To minimize sequence variability in regions of the toehold switch that contribute to secondary structure, the second-generation toehold-switch design (tsgen2, series A) from Pardee et al. was selected for the riboregulator library (25). In this toehold-switch architecture, the target RNA only unwinds six base pairs into the stem, and the top part of the switch is conserved (Supplementary Fig. 1). While other studies have examined complementarity of the entire hairpin stem to the target, we sought to minimize the diversity of RNA hairpins to exclusively examine target-sensor interactions. MATLAB and NUPACK code to design the second-generation toehold-switch architecture may be found in Wu et al. (39). All possible toehold switches across the mCherry RNA target were designed starting from index 21 of the RNA sequence to avoid a putative cryptic ribosome biding site. Of the 655 possible designs, every third index was chosen. If the design featured an in-frame stop codon downstream of the start codon of the toehold switch, the index following the tile with the stop codon was selected. For each toehold-switch variant, the template DNA oligo was designed to contain a 5’ T7 promoter sequence and a common 3’ sequence featuring a 21-nt linker and the first 9 nt of the GFP reporter gene. The GFP variant used in this study is mut3b-GFP with an ASV degradation tag. Cognate 36-nt truncated target transcription templates for each toehold switch featured a 5’ T7 promoter sequence followed by a common stabilizing hairpin and a common 3’ region featuring the first 30 nucleotides (nt) of the T7 terminator. All oligos were ordered as ssDNA from Integrated DNA Technologies.

### Plasmid construction

A two-plasmid system was utilized to transcribe the target RNA in trans to the OFF-state switch RNA on pColE1 and pColA plasmids, respectively. The commercially available pColADuet-1 (EMD Millipore) was used for switch construction and pET15b (EMD Millipore) was used to construct target plasmids. For full length mCherry RNA target expression, the mCherry gene was synthesized and cloned into a pET15b plasmid downstream of a T7 promoter and ribosome binding site (Twist Biosciences).

Plasmid assembly of toehold switches and truncated targets was automated using a Hamilton Microlab NGS STAR platform, with methods encompassing molecular cloning (PCR amplification, Gibson assembly, chemical transformation) and plasmid purification. Template DNA was dispensed from IDT V-bottom storage plates (Corning P-96-450V-B) into a skirted PCR plate (Bio-Rad HSP9601). The robotically assembled common enzyme mix consisted of 2.5 µL ultrapure DNase/RNase-free DI water (Thermo Scientific A57775), 0.5 µL of each 10 µM primer (Integrated DNA Technologies, see Supplemental Table 1), and 5 µL of Q5 2x Master Mix (New England Biolabs M0492L) for each reaction, assembled in a 1.7 mL tube (Genesee Scientific 22-281). Amplification was performed using the on-deck thermal cycler (Inheco) for 35 cycles. Two additional manually prepared 50 µL PCR reactions were performed to linearize the ColE1 and ColA plasmids with vector-specific primers (Supplemental Table 1). PCR was followed by DpnI digestion (New England Biolabs R0176S), column purification (New England Biolabs T1130S), and normalization to 100 ng/µL in ultrapure DNase/RNase-free DI water. Gibson assembly used unpurified insert switch or target PCR products and a custom 2x enzyme mix containing 4 U/µL T5 exonuclease (New England Biolabs M0663S), 25 U/µL Phusion polymerase (New England Biolabs M0530S), 4 U/µL Taq ligase (New England Biolabs M0208L), and 7 ng/µL ET SSB (New England Biolabs M2401S) in Gibson buffer [1 mM dNTPs (New England Biolabs N0446S), 5 mM NAD+ (New England Biolabs B9007S), 500 mM Tris-HCl (VWR 101641-844), 50 mM MgCl₂ (Sigma 63069), 50 mM DTT (Sigma 43816), 25% PEG-8000 (Sigma P2139)]. Each reaction received 1.5 µL of target or switch insert with 0.5 µL of corresponding linearized vector and 2 µL of enzyme mix and was incubated at 50°C for 1 hour. Transformations were performed with in-house prepared chemically competent DH5α *E. coli* (see *Bacterial strains and reagents*), followed by a 1-hour recovery at 37°C in SOC media (Thermo 15544034) in deep-well plates (Thermo AB0859) shaking at 750 rpm. Cells were plated on LB agar (Sigma L3147) with appropriate antibiotics on a rectangular plate (Thermo Scientific 12-566-41) and grown overnight at 37°C. Plasmids were purified with an automated protocol using the Zyppy-96 MagBead Miniprep Kit (Zymo Research D4102). All plasmids were verified by Sanger sequencing.

### Bacterial strains and reagents

*E. coli* DH5α and BL21 (DE3) (New England Biolabs C2987H, C2527H) were used for molecular cloning and toehold switch screening, respectively. Chemically competent cells were prepared following a previously published protocol (40). Cells were streaked on LB agar plates (Sigma L3147), and single colonies were inoculated in LB medium (Sigma L3522) in 14-mL tubes (Corning 352051), and grown overnight 37°C. After 18 hours, cells were diluted 1:100 in fresh medium to a final volume of 500 mL. Cultures were grown to mid-log phase (OD600∼0.4-0.5), chilled on ice for 10 minutes, and pelleted in large centrifuge tubes (Corning CLS430776). Cells were resuspended in 200 mL of ice-cold, 0.22 µm filter-sterilized TBFI [pH 5.2, 30 mM potassium acetate (Sigma P1190), 100 mM rubidium chloride (Sigma R2252), 10 mM calcium chloride dihydrate (Sigma C3381), 50 mM manganese (II) chloride tetrahydrate (Sigma M3634), and 15% (v/v) glycerol (Sigma G6279)], incubated for 15 minutes, pelleted, and resuspended in 10 mL TBFII [pH 6.5, 10 mM MOPS (Sigma M3183), 75 mM calcium chloride dihydrate, 10 mM rubidium chloride, and 15% (v/v) glycerol]. After another 15-minute incubation, cells were aliquoted into 50 µL volumes in pre-chilled PCR plates (Bio-Rad HSP9601), flash-frozen in a dry ice-ethanol bath, and stored at −80°C.

### Flow cytometry

Each toehold switch tile was tested in three conditions: toehold switch with full-length mCherry target (ON-full), toehold switch with truncated target (ON-truncated), or toehold switch with noncognate target (OFF, 36-nt random sequence generated by NUPACK). Each condition was tested in biological triplicate: three colonies transformed with each plasmid combination were inoculated in 500 mL of LB (Sigma L3522) with appropriate antibiotics (50µgmL^−1^ kanamycin, 100 µgmL^−1^ carbenicillin) and grown at 37°C at 750 rpm for 16 hours. 5 µL of overnight culture was then diluted into fresh media with 25µgmL^−1^ kanamycin and 50µgmL^−1^ carbenicillin, grown until OD600 reached 0.2, and induced with 0.25 mM isopropyl β-D-1-thiogalactopyranoside (IPTG, Sigma I6758) for 3 hours. For fluorescence analysis, cultures were diluted 3:1 in phosphate-buffered saline (Sigma P4417) in 384-well plates (Corning 3571). Flow cytometry was performed using a Stratedigm S1000 equipped with an A600 high-throughput auto sampler. At least 10,000 events were collected for each sample and gated based on forward and side scatter profiles (see Supplementary Fig. 2 for gating strategy). Data were analysed using FlowJo^TM^ v10.8 Software (BD Life Sciences). Mean fluorescence values were calculated across three technical replicates for each biological replicate (9 total measurements for each ON- and OFF-state construct).

### Calculations made with ViennaRNA, NUPACK, and RBS Calculator

Thirty-three free energy and structural calculations across the toehold switch were used to assess the average number of nucleotides incorrectly paired at equilibrium: eleven minimum free energy (MFE) values, eleven ideal ensemble defect (IED), and eleven native ensemble defect values (NED). Thermodynamic and structural analyses were performed using a custom Python wrapper for NUPACK 4.0 (27, 41) and previously published python-based ViennaRNA library (28). To extract functionally relevant parameters, each toehold switch RNA sequence was segmented into biologically meaningful regions (toehold domain a, invasion domain b, conserved hairpin, complement b*, linker, and start of GFP) as illustrated in Supplementary Fig.1. MFE was computed for each region using NUPACK and ViennaRNA secondary structures specified in dot-parens-plus notation (where unpaired bases are dots, base pairs are matched paratheses, and strand breaks are indicated by plus signs). Ensemble defect used both the ideal structure (defined below) or the native structure (defined as the structure calculated from MFE) dot-parens representation for each sequence. The ideal structures in run-length-encoded (RLE) dot-parens-plus and DU+ notation are listed in Supplementary Table 2 (41).

To assess the structural accessibility of RNA target regions, MFE and ensemble defect parameters were calculated for 36-nt binding sites and surrounding flanking sequences. For each candidate binding site, a widow spanning the target site and 10, 25, 50, or 100-nt flanks was extracted from the full-length RNA transcript. MFE, ideal structure defect, and native structure defect were calculated, with the ideal structure defined as a completely unpaired RNA sequence. To quantify translation-related features that may influence target availability, the average codon usage fractions across regions surrounding the RNA target site were calculated. Each 36-nt target sequence was located and aligned to the nearest in-frame codon. The average codon fractions were computed for: the first two in-frame codons (the two codons in the 3’ hairpin stem of the toehold switch b* region), the entire 36-nt binding site, 4-10 codons upstream of the site within the target coding sequence, and 4-10 codons downstream within the target coding sequence.

Base pairing probabilities were obtained from the total partition function computed by summing over all possible secondary structures weighted by their Boltzmann factors using NUPACK and ViennaRNA. For each base, the pair probabilities greater than 0.01 were selected and saved. Base-pairing interactions across the full RNA target predicted from pairwise probability data were visualized using an arc plot generated by NetworkX (https://github.com/networkx/networkx).

RBS Calculator predictions were made by the most recent publicly available version (42–45). For each switch, the maximum translation initiation rate (TIR) of the on-target start codon was predicted for either the ON or OFF state as specified by the “Switch ON-GFP” or “Switch OFF-GFP” structures above.

### Frequency logos

To identify enriched sequence-level motifs associated with toehold-switch performance, we applied kpLogo to compare the best and worst performing variants (46). All switches were ranked based on the five experimental ON, OFF and fold change functional values. The top and bottom 10% of each distribution were selected and used as input for kpLogo to reveal statistically overrepresented motifs at specific nucleotide positions. Default settings were used, and analyses were conducted in nucleotide mode and with input sequences ranked based on experimental data.

### Machine learning model

To evaluate and predict the performance of RNA sensors based on sequence-derived thermodynamic and structural parameters, we implemented a supervised classification pipeline using Partial Least Squares Discriminant Analysis (PLS-DA). Raw data were loaded from a CSV file containing all five experimental performance metrics and 68 associated input features derived from RNA folding predictions and structural calculations. Feature columns were standardized using a z-score normalization to ensure each variable contributed equally to downstream modeling. For classification, each target metric was binarized into “High” and “Low” classes by thresholding at the 65th percentile of each metric’s raw distribution. Labels were encoded using a binary integer mapping.

To reduce dimensionality and mitigate overfitting, we performed Recursive Feature Elimination (RFE) with a logistic regression estimator, retaining the top 20 features for each classification task. RFE was chosen to prioritize model interpretability while preserving performance, and selected features varied depending on the target metric. The reduced feature sets were then used to train PLS models across a range of 1–20 components for each independent experimental metric.

To determine the optimal number of latent components, a 5-fold cross-validation strategy was implemented. For each number of components, we used the latent variables from the PLS model as input to a logistic regression classifier and recorded mean classification accuracy. The number of components that maximized cross-validation accuracy was selected for final model construction, and the cross-validation error was reported as one minus the best mean accuracy to ensure model generalizability and avoid overfitting due to excessive latent dimensions (Supplementary Fig. 3). Each finalized PLS-DA model was trained using the reduced training data, and latent features from the test set were used to evaluate classification performance using accuracy, confusion matrices, and ROC curves with area under the curve (AUC) as the primary metric.

A key parameter influencing model performance was the threshold used to binarize values for classification. Instead of applying a fixed cutoff, we used a percentile-based approach, defining “High” performers as those above the 65th percentile of each metric’s raw distribution and “Low” performers as those below. The class balancing strategy significantly impacted AUROC and other PLS-DA metrics such as precision, recall, and F1 score for each of the five empirically determined metrics. The rationale and performance trade-offs underlying this percentile-based thresholding are presented in Supplementary Fig. 3.

### Toehold-VISTA

To facilitate the adoption of Toehold-VISTA, code has been made available as a GitHub repository (https://github.com/AlexGreenLab/vista). All modules are accessible as Jupyter notebook code, which can be used for any new Toehold-VISTA analysis. Toehold-VISTA was developed to automate tiling, thermodynamic modelling, and machine-learning guided scoring of toehold switches. The Toehold-VISTA code was built upon the canonical toehold switch architecture, wherein the 5’ hairpin stem, loop, and 3’ hairpin stem was fixed using the sequence 5 ′ – GUUAUAGUUAUGAACAGAGGAGACAUAACAUGAAC – 3 ′. A 36-nt trigger-binding domain was implemented, which invades the switch stem by 6 nt. Designs were initiated by uploading a CSV file containing the RNA target name, sequence, desired temperature for the NUPACK model, output/reporter name, and the sequence of the reporter. In each instance of the design function, toehold switches were designed to target either a truncated or full RNA sequence, as specified.

For each RNA input, Toehold-VISTA generated all possible 36-nt target sequences tiled across the transcript and their cognate switch. Candidate toehold switches then underwent filtering and sequence validity checks. Toehold switches were excluded if they contained premature in-frame stop codons prior to the output gene. Valid designs were scored using the previously trained PLS-DA model parameters. Designs were ranked based on their predicted ON/OFF performance using the corresponding model parameters. Final rankings and scores were exported, along with a file of putative high-performing toehold switches and their targets.

### Statistical analysis and reproducibility

All experimental characterization of toehold switches were repeated in three biological replicates with three technical replicates, for a total of nine experimental measurements for each toehold switch. All machine learning results, including reported cross-validation performance metrics and unreported model tuning and feature selection steps, were independently replicated to ensure reproducibility. This includes rerunning each PLS-DA model, linear regression model, and PCA analysis using separate initialization and training iterations.

## RESULTS

### Library synthesis and screening

Current design paradigms in RNA synthetic biology primarily focus on intrinsic RNA sensor sequence and structural properties while often placing less emphasis the structural complexity and accessibility of the RNA target itself. While recent attempts have been made to apply various machine learning and deep learning techniques to RNA sensor design, all omit consideration of the target RNA’s structure, limiting predictive power and design success (34–36). To improve our understanding of trans RNA-RNA interactions and our ability to design RNA-based sensors, we constructed an experimental system that allowed us to quantify the functional consequences of targeting distinct regions across a structured RNA molecule using the toehold switch. The toehold switch contains a cis-repressing RNA hairpin that blocks the ribosome binding site (RBS) and start codon, inhibiting translation of a downstream reporter. Upon activation by a trans-acting target RNA, the target RNA binds to the exposed toehold sequence of the switch, relieving repression and enabling downstream gene translation (2).

A comprehensive library of toehold switches was tiled along the length of an mCherry RNA transcript at 3-nt intervals, yielding a total of ∼200 toehold switches (Figure 1A). Each variant was designed using the second-generation toehold switch scaffold (47), preserving the conserved structural backbone while varying the 36-nt trigger-binding domain. Each switch was screened in *E. coli* BL21(DE3) using a two-plasmid system under three conditions: an OFF state with a noncognate trigger, an ON state with a truncated 36-nt cognate target, and an ON state with the full-length mCherry transcript as the cognate target. We included truncated targets containing only the 36-nt trigger region to decouple sensor activation from long-range RNA folding effects and to establish a baseline for hybridization efficiency in the absence of structural occlusion. Flow cytometry data revealed a broad distribution of ON-state GFP fluorescence across both truncated and full-length targets (Figure 1C, Supplementary Fig. 2), confirming that a range of activation efficiencies was observed across the library. Notably, toehold switches activated by full-length targets exhibited significantly reduced fluorescence and fold change relative to truncated targets (p<0.0001, paired two-tailed t test), consistent with a role for native RNA structure in limiting sensor-target hybridization and activation (Figure 1D). These results highlight the relationship between target RNA structure and toehold switch performance and indicate that thermodynamic or structural properties of the full-length target RNA impact device function.

**Figure 1.**
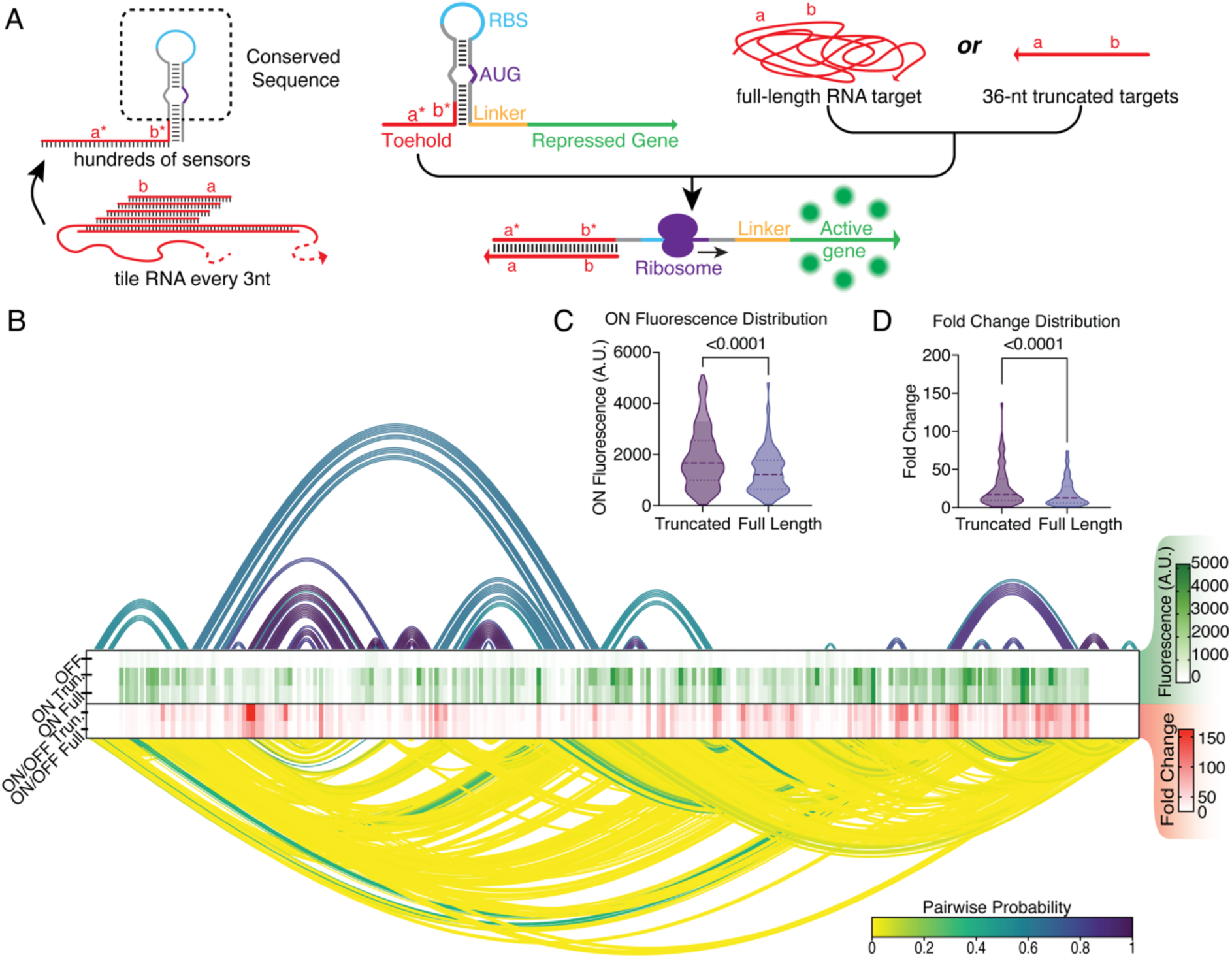
Screening of toehold switches across the mCherry RNA transcript. (A) Schematic of the toehold switch library design, in which switches were tiled at 3-nucleotide intervals along the mCherry transcript using a conserved hairpin scaffold sequence and varying 36-nt binding domains. (B) Arc diagram of pairwise probabilities with most probable (top) and least probable (bottom) interactions plotted by increasing nucleotide index from left to right. Heat map of OFF-state, ON-truncated, and ON-full RNA GFP fluorescence and fold change data are aligned to each respective index. (C) Quantification of ON state GFP fluorescence for truncated and full-length RNA transcripts (p < 0.0001, two-tailed t test). (D) Fold-change distribution for truncated and full-length activation of toehold switches (p < 0.0001, two-tailed t test). For violin plots, the horizontal dashed line represents the median and dotted lines are at 25^th^ and 75^th^ percentiles.

### Mechanistic switch evaluation based on RNA secondary structure predictions

To better understand the biophysical mechanisms that underlie toehold switch performance, we analyzed how predicted secondary structure features of the switch RNA relate to functional outputs. Approaches to design RNA-based synthetic biology tools largely rely on thermodynamic models of RNA secondary structure (2, 39). Building on these established methods, we used the NUPACK (41) and ViennaRNA (28) software suites to calculate 33 thermodynamic and structural parameters—encompassing minimum free energy (MFE), ideal ensemble defect (IED), and native ensemble defect (NED)—for eleven biologically meaningful regions of each toehold switch. We then evaluated how these parameters correlated with the five key empirically determined metrics (Figure 2A, Supplementary Fig. 4A).

**Figure 2.**
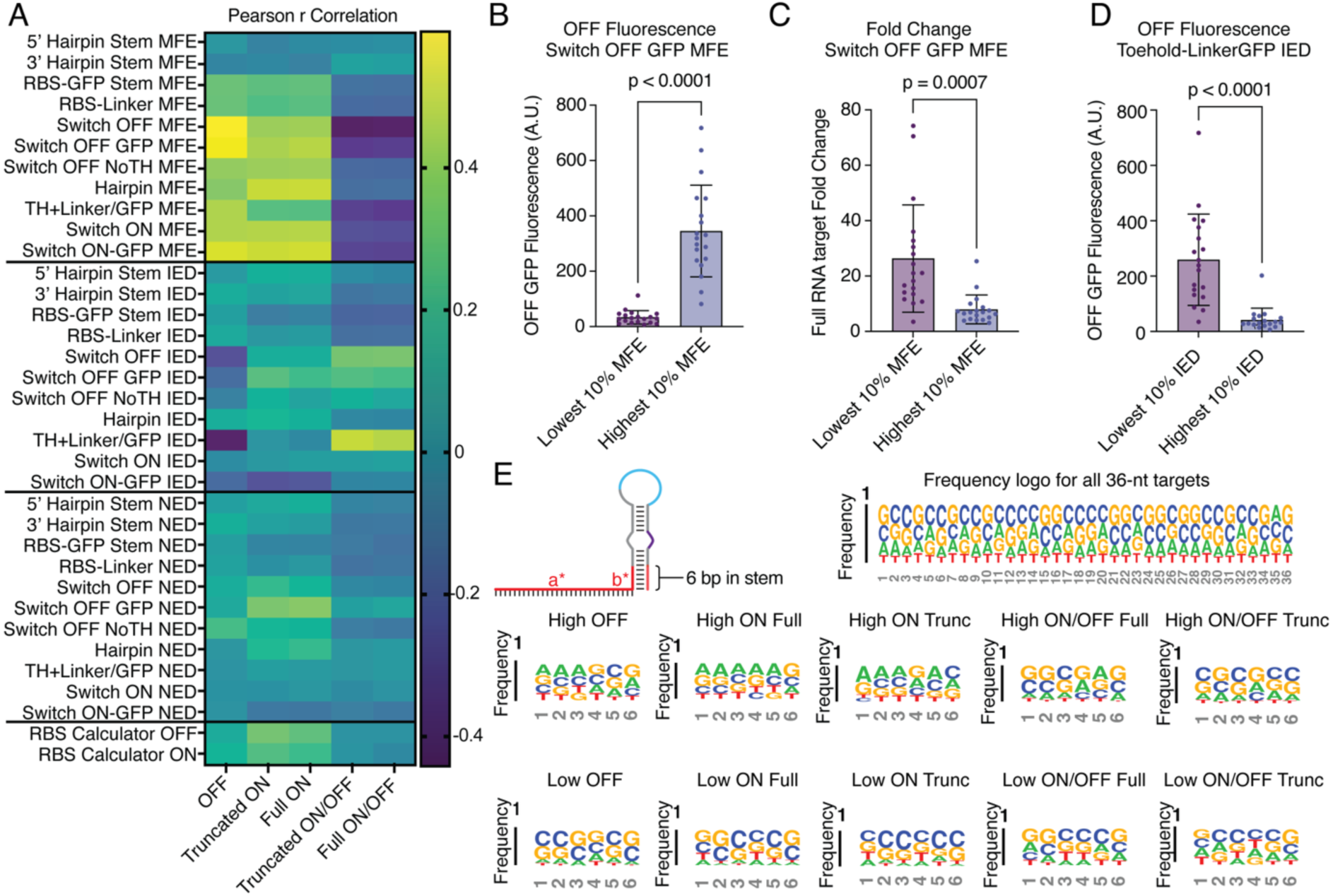
Rational feature determinants of toehold switch performance. (A) The Pearson correlation (max(|r|) = 0.5) between 33 thermodynamic and structural features of toehold switches, RBS calculator v2.1 outputs, and experimentally determined values. (B) OFF-state GFP fluorescence stratified by lowest and highest 10% (n=19) NUPACK predicted OFF-state-GFP MFE (p < 0.0001 two-tailed t test). (C) Full RNA target fold change stratified by lowest and highest 10% (n = 19) NUPACK predicted OFF-state-GFP MFE (p = 0.0007 two-tailed t test). (D) OFF-state GFP fluorescence stratified by lowest and highest 10% (n=19) NUPACK calculated ideal ensemble defect for toehold-linker structure. (E) Sequence logos for all 36-nt RNA target (top) and for the first 6 nt at the base of the 3’ end of the toehold switch hairpin stem discovered to be disproportionately represented in various functional groups. For all bar plots, horizontal line indicates the median and whiskers indicate the standard deviation (SD).

The most pronounced relationships were observed in the OFF state, where MFE values of the ribosome binding site (RBS) RBS-GFP stem, RBS-Linker region, and overall switch OFF state showed the strongest positive correlations (Pearson r > 0.45) with background fluorescence. This suggests that less thermodynamically favorable OFF structures, reflected by higher (less negative) MFE values fail to sequester the RBS and start codon effectively leading to undesired leaky expression. A similar trend was observed with ON-state fluorescence, though weaker in magnitude, with MFE correlating positively with ON GFP fluorescence.

To further validate these trends, we stratified switches by their predicted MFE and compared empirical OFF fluorescence. Switches in the lowest 10% of OFF-state GFP MFE exhibited significantly reduced background fluorescence compared to the top 10% (p<0.0001, two-tailed t test), confirming that thermodynamic free energy of the OFF hairpin is a major determinant of leaky expression (Figure 2B). Visualization of representative structures highlights this difference: the lowest OFF switch (index 662, MFE ΔG = –54.25 kcal/mol) exhibited strong pairing between the linker and toehold region, while the highest OFF switch (index 592, MFE ΔG = –27.33 kcal/mol) formed weaker structures with exposed RBS and start codon (Supplementary Fig. 4B, D). Importantly, when comparing fold change across the extremes of MFE values, lower MFE also corresponded to significantly higher fold change (p = 0.0007, two-tailed t test; Figure 2D). However, switches with the lowest 10% predicted ideal ensemble defect (IED) values for the toehold-linker-GFP region also showed markedly higher OFF-state fluorescence compared to those with the highest 10% ideal ensemble defect values (p < 0.0001, two-tailed t test; Figure 2D, Supplementary Fig. 4C). Therefore, toehold regions that are largely single stranded may provide greater accessibility for target sensing but are more susceptible to leaky OFF state expression (Supplementary Fig. 4E).

To examine how sequence content may contribute to these structural trends, we analyzed the nucleotide base composition of the first 6 nucleotides of the 3’ hairpin stem, which correspond to the 5′ end of the target, across switches ranked by ON/OFF performance. Sequence logos revealed that high-performing switches—those with high fold change and low OFF fluorescence—were enriched for balanced GC and AU content, likely stabilizing the stem (Figure 2E). In contrast, switches with high OFF fluorescence showed strong enrichment for adenines in this region, potentially forming weak stems that fail to properly repress translation in the absence of target RNA. Similarly, switches with low ON-state fluorescence (for either truncated or full targets) displayed GC enrichment, suggesting that overly rigid stems may also impede effective target-triggered unfolding and activation.

To assess whether more comprehensive models of translation initiation could improve predictive power, we applied the RBS Calculator to estimate maximum translation initiation rates for both ON and OFF states (43–45, 48). While modest correlations were observed between predicted ON initiation rates and experimental ON fluorescence (Pearson r = 0.27-0.34), and weaker correlations for the OFF state (Pearson r = 0.12-0.17), these values were comparable to or lower than those from simpler thermodynamic features (Figure 2A), suggesting that existing RBS-based models may not fully capture the structural and contextual complexities of toehold switch regulation and activation.

Taken together, these results highlight the limitations of existing thermodynamic and kinetic models in fully capturing toehold switch behavior. While some structural parameters show modest correlations with function, many fail to consistently predict sensor performance across conditions, possibly due to unaccounted factors in the cellular environment resulting in inaccuracies in energetic folding models developed from in vitro measurements. Crucially, these findings underscore the importance of considering not just the sensor’s structure, but also the structure and accessibility of its RNA target.

### Target structure-guided analysis using RNA folding models

To better understand how RNA secondary structure of the target transcript influences toehold switch activity, we first developed a contextual modeling framework that quantifies base pairing probabilities across different regions of the truncated 36-nt target (Figure 3A). Specifically, we calculated average base-pairing probabilities from ViennaRNA-generated pair-probability matrices for several biologically relevant subregions of the 36-nt target site: the first 3, 6, and 18-nt from the 5′ end, which are designed to initiate hairpin opening in the toehold switch; and the first 3, 6, and 18-nt from the 3′ end, which theoretically nucleate target-sensor binding. Additionally, we computed the average pair probability across the full 36-nt sensor binding site. Across these features, we observed modest correlations with experimental ON values, most notably with average pair probabilities over the 5′ 18-nt (Pearson r = −0.27) and full 36-nt region (Pearson r = −0.38), suggesting that reduced structural constraints in the target correlates with higher activation (Figure 3B, Supplementary Fig. 5A). When we stratified the highest and lowest pair probabilities for the entire 36-nt truncated target, we found a significant decrease in ON-state GFP fluorescence (Figure 3C). Interestingly, weak correlations were also observed between 3′ end pairing of the target and ON/OFF ratios (Pearson r = −0.33), potentially reflecting kinetic effects during toehold initiation.

**Figure 3.**
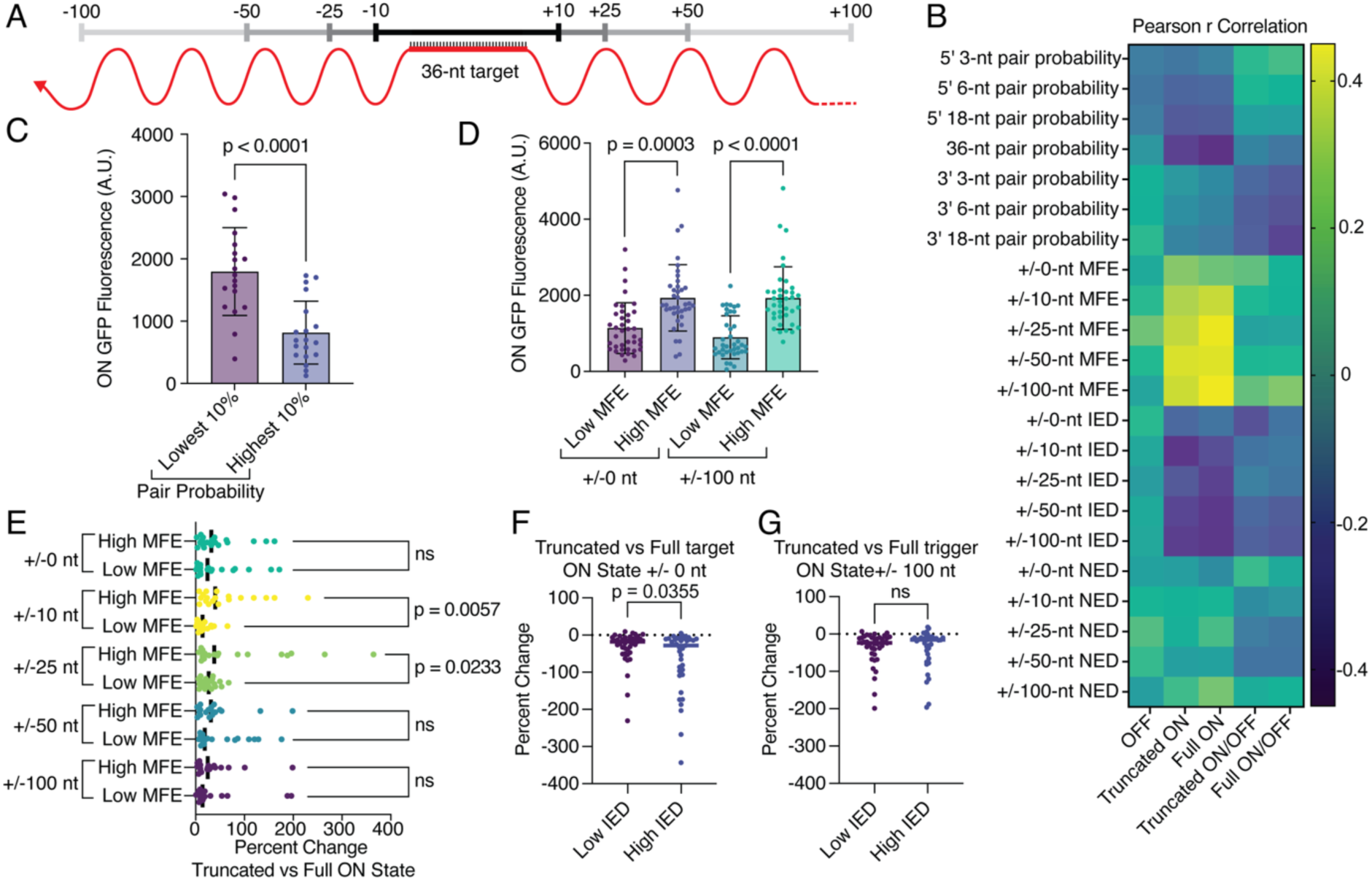
Influence of target RNA structure and context on toehold switch activation. (A) Schematic of contextual modelling used to compute rational metrics for the 36-nt sensor binding site and surrounding sequences with variable flanking lengths. (B) Pearson correlation (max(|r|) = 0.43) across 22 thermodynamic and structural features of the target RNA and experimentally determined values. (C) ON-state GFP fluorescence stratified by the lowest and highest 10% (n=19) average pair probabilities for the 36-nt target window (p < 0.0001, two-tailed t test). (D) ON-state GFP fluorescence stratified by the lowest and highest 20% (n=38) minimum free energy (MFE) for ±0 or ±100 nt flanks (p = 0.0003 and p < 0.0001, respectively; two-tailed t test). (E) Percent change in measured ON-state GFP fluorescence between truncated target and each full-length RNA target flanking length, stratified by the lowest and highest 10% (n=19) MFE (two-tailed t tests). (F, G) Percent change in measured ON-state GFP fluorescence between truncated and full-length target with no flank (F) or 100-nt flanks (G), stratified by the lowest and highest 20% (n=38) ideal ensemble defect (p = 0.0355, ns two-tailed t test). Bars in (C, D) represent the median and whiskers represent standard deviation (SD). Lines in (E, F, G) represent the median.

We next developed a modeling approach to capture additional contextual influences by incorporating varying lengths of target-flanking sequence windows (Figure 3A). Minimum free energy (MFE), ensemble defect metrics, and codon usage parameters were calculated for the 36-nt sensor binding site, as well as for extended windows of ±10, ±25, ±50, and ±100-nt adjacent to the sensor binding site. Across these structural and thermodynamic features, we observed that MFE consistently correlated more strongly with full-length RNA target ON-state fluorescence than with the truncated RNA target (Figure 3B). Notably, the correlations peaked with a ±25 and ±100-nt window (Pearson r = 0.41, 0.43 respectively), suggesting that more extended structural context enhances predictive power. In contrast, ensemble defect metrics such as ideal ensemble defect showed negative correlations with ON state fluorescence, particularly for wider windows (e.g., Pearson r = −0.36 at ±100 nt), reflecting that more structured regions surrounding the sensor binding site reduce activation efficiency. These findings suggest that sensor binding site availability, modulated by local and global folding of the target, is a critical determinant of switch performance.

To test this further, we stratified switches by MFE and compared GFP fluorescence outputs. While sensor binding sites with high MFE (i.e., less structured) consistently led to higher ON-state fluorescence, the magnitude of this difference increased dramatically when flanking regions were included in the folding model. Specifically, the GFP signal difference between high and low MFE sensor binding sites increased by 362.6% for ±100-nt windows compared to only 141.9% with no flanking sequence (Figure 3D), emphasizing the importance of accounting for broader transcript context when modeling sensor-target interactions.

We next assessed whether local or distal secondary structure differentially impacted sensor activation through comparisons between the full-length targets and truncated targets. When stratifying toehold switches based on MFE and comparing the relative fluorescence change between truncated and full-length RNA triggers, we found that only shorter flanking windows (±10-nt and ±25-nt) showed statistically significant differences (p = 0.0057 and p = 0.0233, two-tailed t test, respectively; Figure 3E). Larger windows (±50-nt and ±100-nt) did not show significant differences in ON-state fluorescence, suggesting that proximal structural interactions near the binding site more directly modulate switch accessibility and activation.

To probe this further, we examined ensemble defect metrics. Stratifying the lowest and highest 20% ideal ensemble defect values revealed a significant drop in ON-state fluorescence for structured targets (high ideal ensemble defect) only in the 0-nt flank model (p = 0.0383, two-tailed t test), but not for ±100-nt contexts (Figure 3F, G). Therefore, proximal occlusion of the binding site rather than global folding of the full transcript more strongly determines accessibility and functional response. Together, these analyses highlight the importance of local RNA secondary structure and thermodynamic accessibility in determining the functional output of toehold switches. Our findings suggest that proximal target structure—rather than distal or global folding—is a dominant factor influencing toehold accessibility.

### Target structure-guided analysis using RNA folding models

Recognizing that transcription and translation are coupled in bacteria—and that ribosome binding may reshape RNA folding—we next investigated whether codon composition near the target site influences toehold switch behavior (49, 50). Several codon usage features, especially those in the first two codons of the sensor binding site, correlated positively with ON state fluorescence (Figure 4A, Supplementary Fig. 6A). When stratifying switches by codon fraction, we found that sequences containing more frequently used codons in the first two positions downstream of the AUG (the 3’ base of the hairpin stem) had significantly higher ON state fluorescence in both the truncated (p = 0.0213, two-tailed t test) and full RNA context (p = 0.0322, two-tailed t test), suggesting that less ribosomal engagement at these sites may enhance switch activation, potentially by leading to more efficient translation (Figure 4B, C, Supplementary Fig. 6B), as has been examined in previous studies (37, 38).

**Figure 4.**
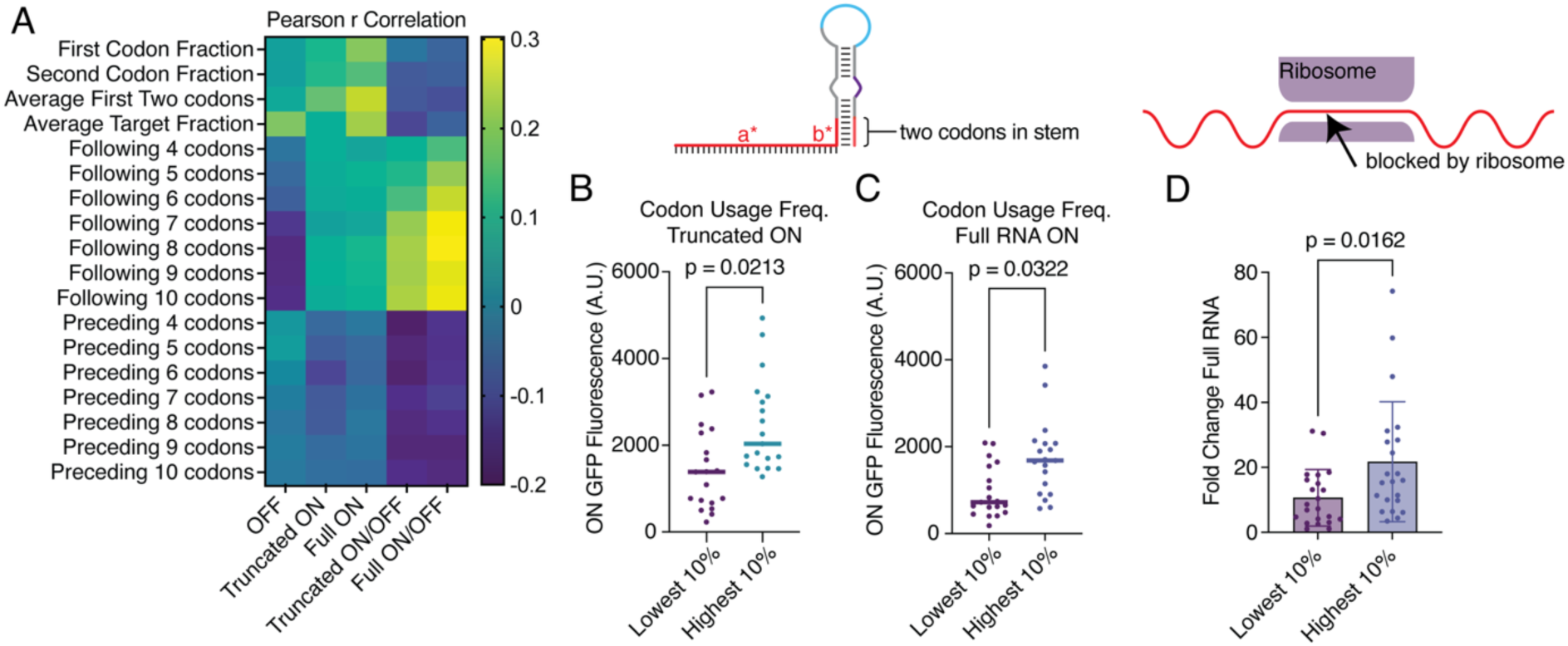
Codon fraction modulates toehold switch activation. (A) Pearson correlation (max(|r|) = 0.3) between 18 codon fraction calculations and experimentally determined values. Comparison between switches with highest and lowest 10% (n=19) codon usage in the variable two codons at the base of the hairpin stem for truncated target (B) and full-length RNA target (C) (p = 0.0213 and p = 0.0322, respectively; two-tailed t test). (D) Full RNA fold change stratified by the lowest and highest 10% (n=19) average codon fraction for the 7 codons following toehold switch target sites (p = 0.0162, two-tailed t test). Lines in (B, C) represent the median. Bars in (D) represent the median and whiskers represent standard deviation (SD).

We also assessed the average codon fractions for various sites surrounding the sensor binding site. We hypothesized that lower codon usage proximal to a sensor binding site may result in ribosome stalling and therefore lower target availability (Figure 4D). This is supported by the understanding that ribosomes progress more slowly through codons that correspond to less abundant tRNAs, which can lead to transient pausing or slowing of translation (51). Such ribosome pausing may arise from biophysical constraints, such as slower accommodation of rare tRNAs in the A-site of the ribosome, or kinetic barriers to peptide bond formation when uncommon codons are present (52). Given that the ribosome footprint typically spans ∼9–10 codons on the mRNA and that the E, P, and A sites—which mediate tRNA binding and peptide elongation—correspond to positions 4, 5, and 6 within this footprint, the presence of slow-translating codons just downstream of a sensor binding site could position the ribosome such that the target region remains covered (53). This prolonged blockage may frustrate toehold switch binding and activation, resulting in the observed lower fold changes when targets are positioned around regions of lower codon usage. We found correlations between each region and experimentally determined values, with the strongest correlation between the average 8 codons following a sensor binding site and the fold change of the full RNA target (Pearson r = 0.30).

Codon-level effects highlight the relevance of translation-linked remodeling of mRNA during switch activation. While these rational metrics provide mechanistic insights into toehold switch performance, their limited predictive power—across both thermodynamic and structural features of switches and their RNA targets—underscores the need for a more complex model. To address this gap, we sought to exploit machine learning as a framework to capture complex multivariate patters and improve functional prediction of toehold switch performance.

### Improved toehold switch functional prediction by partial least squared discriminant analysis

Given the complex interplay between thermodynamic and structural context-dependent features, we employed Partial Least Squares Discriminant Analysis (PLS-DA) to capture these multivariate relationships. PLS-DA was selected as it allowed for a supervised machine learning approach capable of reducing the dimensionality of the data while preserving class separation. PLS-DA is well-suited for high-dimensional, multicollinear datasets with limited sample sizes, as it performs simultaneous dimensionality reduction and regression while maximizing class separation in the latent variable space. In this way, we aimed to robustly distinguish high- and low-performing toehold switches while enhancing interpretability through latent variables and feature loadings, enabling better insight into the parameters that most strongly influence switch function than other black-box deep learning models.

First, we used recursive feature elimination (RFE) with logistic regression to select the top 20 most-informative features from the initial set of 66 calculated thermodynamic, structure, and sequence-derived parameters (Figure 5A). Cross validation was then used to determine the optimal number of latent variables for each PLS-DA model (Figure 5B). Across all five experimental readouts, classification accuracy improved significantly within 5 components. The highest accuracy was observed for both truncated and full-length RNA fold-change models, achieving mean cross-validation accuracies above 90% with 7 components. The selected components were then used in each individual PLS-DA model. Strong classification performance was achieved across all five models (Figure 5C), with area under the curve (AUC) values of 0.96 (ON/OFF Truncated) and 0.98 (ON/OFF Full RNA) indicating excellent sensitivity and specificity. These metrics were further supported by high precision, recall, and F1 scores (Supplementary Fig. 3). Importantly, we found that model performance was sensitive to the binary threshold used to classify fold change values. A detailed threshold optimization analysis revealed that a threshold of 0.65 maximized the AUC for the ON/OFF Full RNA model balancing a precision value of 1.00, recall of 0.87, and F1 score of 0.93 at this cutoff (Supplementary Fig. 3 A-E).

**Figure 5.**
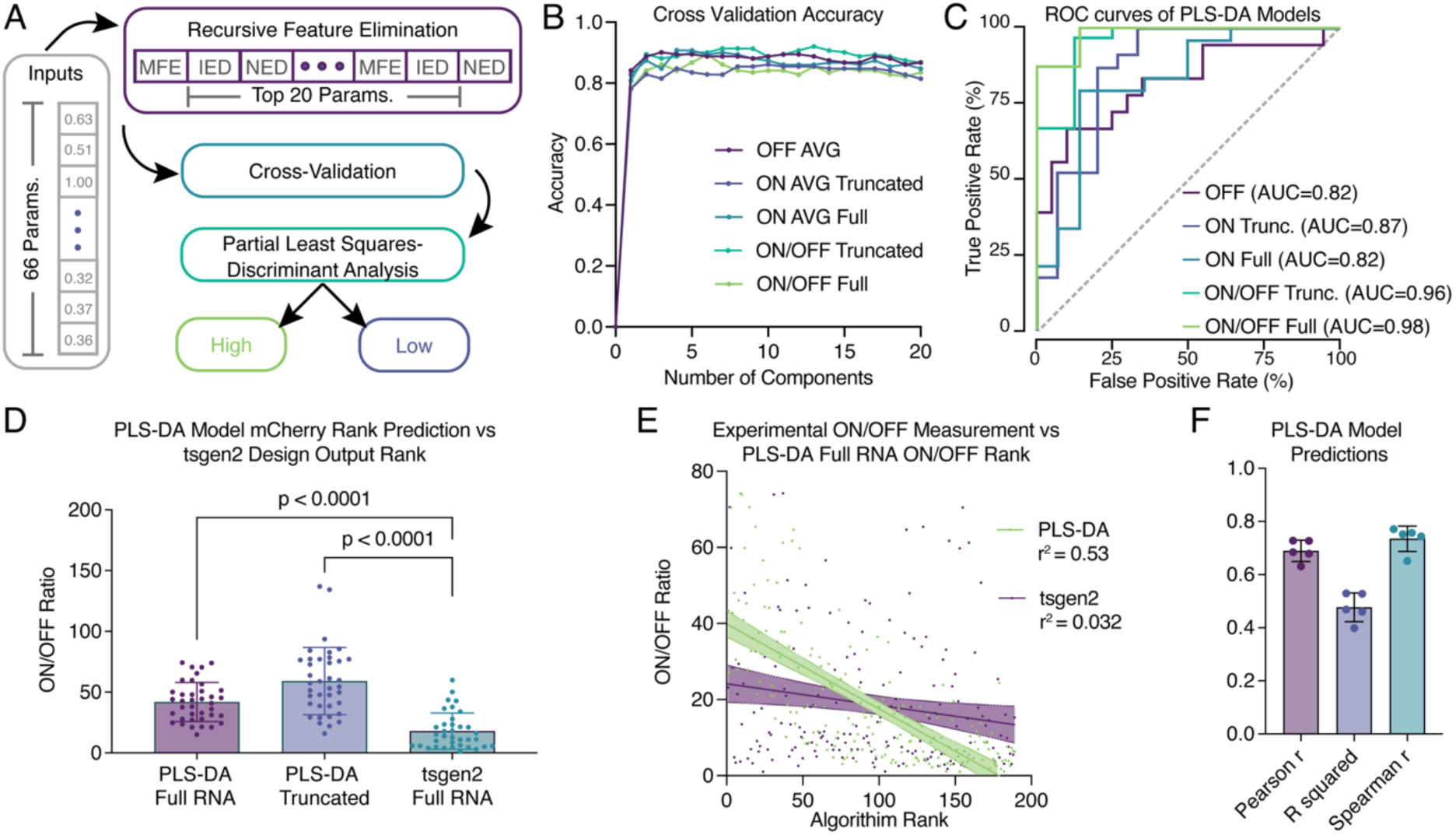
Partial least squares discriminant analysis (PLS-DA) enables robust prediction of toehold switch function. (A) Top 20 predictive features were selected from 66 rational parameter inputs via recursive feature elimination. (B) Cross-validation accuracy for selection of the optimal number of components for PLS-DA models across experimental outputs. (C) Receiver operating characteristic (ROC) curves for each model. (D) Comparison of ON/OFF fold change for the top 20% of switches (n=19) ranked by PLS-DA for full RNA and truncated targets, and tsgen2 rankings. Bars represent median and whiskers represent SD. (E) Experimental ON/OFF fold change values for full RNA target and PLS-DA algorithm rank. Linear regression is plotted with 95% confidence interval. (F) PLS-DA model predictions correlations with experimentally determined values. Bars represent median and whiskers represent SD.

Feature contributions to each latent variable were visualized using PLS loadings, enabling interpretation of how individual sequence and structure features impacted sensor classification (Supplementary Fig. 7A, B). These loading values were plotted for the first latent variable to highlight the most influential features for each model. Using PLS-DA loading plots, we identified common and model-specific features most predictive of toehold switch function. Across all models, Switch ON MFE, ON-GFP MFE, and OFF MFE toehold switch sequence regions consistently had the highest loading values (0.48, 0.53, and 0.50, respectively), suggesting that the free energy of the activated and inactive state of the toehold switch is a dominant driver of toehold function. Ensemble defect values emerged as important in several models, especially the OFF-state and truncated-target models. Specifically, the toehold/linker ideal ensemble defect and switch OFF ideal ensemble defect (loading values of 0.45 and 0.31, respectively) attributed most strongly to the truncated fold change. Thermodynamic and structural parameters of the target were reflected strongly in PLS-DA models for the full RNA target ON state, with target ideal ensemble defect values for the flanks ± 0, 10, 25, and 50-nts reflecting negative loadings. Since the ideal structure in our analysis is defined as unpaired, the negative loading indicates that more base pairing negatively impacts performance: as the ideal ensemble defect increases, the model predicts worse performance. Therefore, in regions with reduced accessibility, hybridization is impeded, resulting in lower switch activation and aligning with the mechanistic requirement for an unstructured target to efficiently invade and displace the switch hairpin.

We next evaluated how well the full-length RNA fold-change PLS-DA model ranked toehold switch candidates as compared to tsgen2, the current standard of toehold-switch design (25, 39). The top 20% of switches identified by the PLS-DA model exhibited significantly higher ON/OFF ratios than those ranked highest by tsgen2 (p=0.0001, two-tailed t-test; Figure 5D), for both full-length and truncated RNA targets. When training the model using only truncated ON/OFF data, the same trend was observed (Supplementary Fig. 7C). The PLS-DA ranking achieved much stronger correlation with actual experimental fold change than tsgen2: the full RNA target ON/OFF PLS-DA model achieved a value of r^2^ = 0.53 (Figure 5E), compared to just r^2^ = 0.032 for tsgen2. The results were consistent across the four other trained models (Supplementary Fig. 7C), demonstrating that PLS-DA captures predictive features that tsgen2 fails to incorporate.

### Toehold-VISTA enables improved sensor design for an unseen RNA target

To demonstrate the practical application of our PLS-DA framework, we integrated it into a pipeline for sensor design and ranking. The workflow, termed Toehold-VISTA, begins by generating all toehold switch candidates across a given RNA target, calculating 66 mechanistic and thermodynamic features per design. From this, the top 20 predictive features—identified via recursive feature elimination in the trained PLS-DA model—are transformed into latent variables using PLS loadings. These are then scored using logistic regression coefficients to assign each candidate a probability of belonging to a high- or low-performing class. The final VISTA output ranks the switches accordingly (Figure 6A).

**Figure 6.**
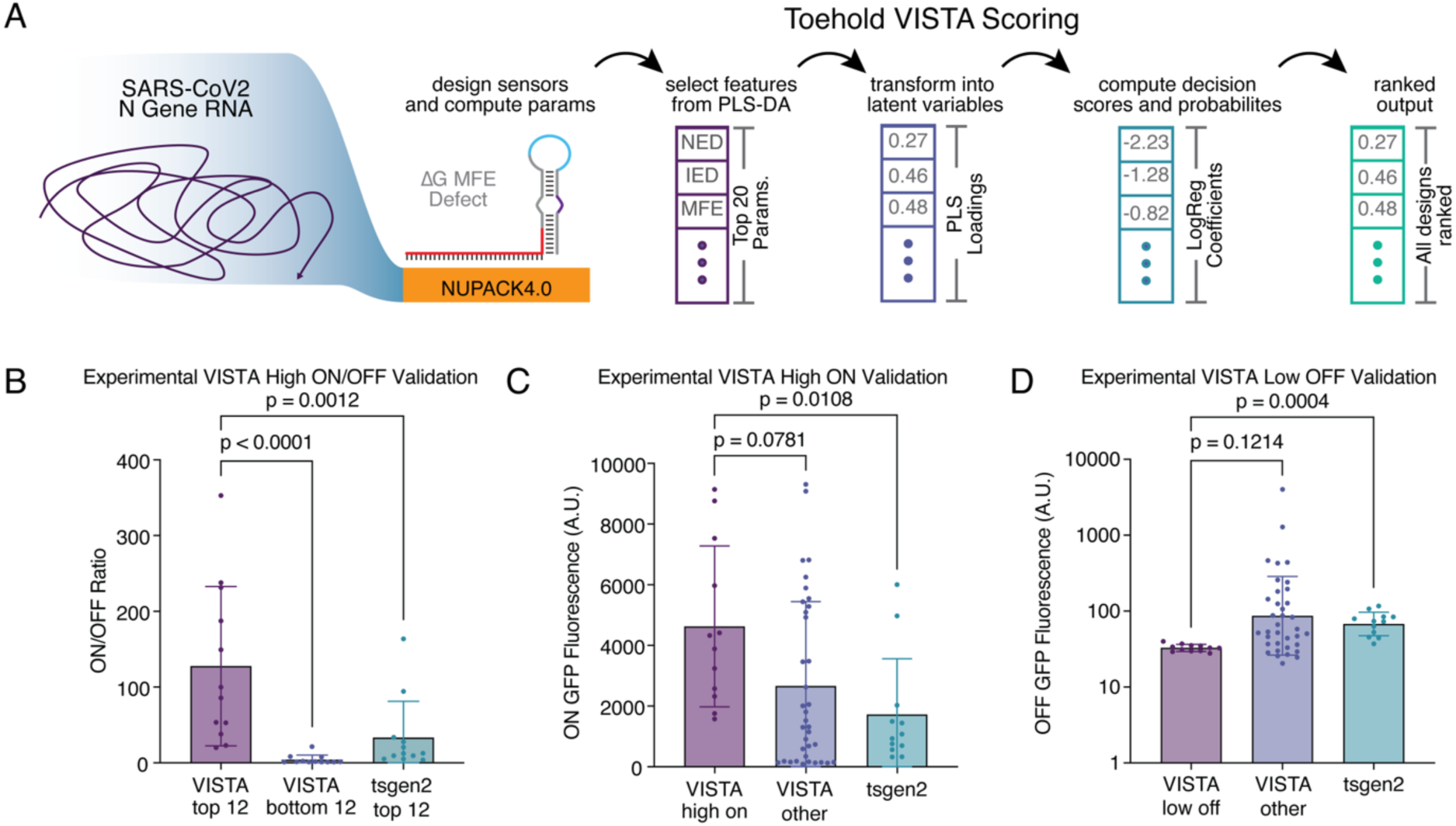
Toehold-VISTA enables accurate functional prediction for a new RNA target. (A) Schematic of the VISTA pipeline: all possible toehold switch candidates are designed with NUPACK and rational parameters are calculated. The top 20 thermodynamic and structural parameters are chosen and the PLS loadings are used to transform data into latent variables. Logistic regression coefficients are finally used to compute probabilities that each switch belongs in the “high performance” class, before returning a ranked output of all possible toehold switches. (B) ON/OFF fold-change data for experimental validation of the top 12 and bottom 12 VISTA-scored toehold switches and top 12 tsgen2-designed toehold switches (comparisons made by one-way ANOVA). (C) ON GFP fluorescence data for 12 VISTA-scored toehold switches for high ON-state fluorescence, all other VISTA-designed constructs (n=36) and top 12 tsgen2 designed switches. (D) OFF GFP fluorescence data for 12 VISTA-scored toehold switches for low OFF-state fluorescence, all other VISTA designed constructs (n=36), and top 12 tsgen2-designed switches. Comparisons in (C, D) made with Dunnett’s T3 multiple comparisons test. Bars in (B, C, D) represent the median and whiskers indicate the SD.

We validated this approach experimentally by applying Toehold-VISTA to a new, unseen RNA target: the SARS-CoV-2 nucleocapsid (N) gene. We first selected the top 12 highest-ranked and bottom 12 lowest-ranked switches predicted by VISTA (using the ON/OFF full RNA target model) and compared their experimental ON/OFF fold-change performance. The top VISTA designs demonstrated significantly higher ON/OFF ratios than both the lowest VISTA-ranked switches (p < 0.0001, two-tailed t test) and the top 12 switches ranked by the prior state-of-the-art tool, tsgen2 (p = 0.0012). This trend held across all 12 pairs, with VISTA high-performing switches exhibiting consistently superior fold change (Figure 5B).

Next, we tested whether VISTA could predict high-ON-state switches using the full-RNA ON fluorescence model (Figure 5C). The 12 switches with the highest predicted ON values were experimentally validated and compared to the entire VISTA pool as well as the top 12 tsgen2-ranked switches. VISTA high-ON designs were significantly higher than the tsgen2 top set (p = 0.0108, two-tailed t test), achieving maximal individual fluorescence values as high as 9140.7 compared to 1075.9 for the top tsgen2 design. We also evaluated the VISTA OFF fluorescence model by selecting the 12 switches predicted to have the lowest OFF state. Although the group with switches selected to have the lowest off state did not reach significance compared to the rest of the VISTA switch pool (p = 0.1214, two-tailed t test), it showed significantly reduced OFF fluorescence relative to tsgen2 (p = 0.0004, two-tailed t test). Notably, this group maintained consistent OFF values for most switches, while tsgen2 exhibited higher baseline OFF levels (Figure 5D).

Together, these experiments validate the VISTA pipeline as a powerful design tool that leverages machine learning to accurately predict and rank toehold switch performance. VISTA significantly outperforms existing tools like tsgen2 in both ON/OFF discrimination and ON-state prediction, highlighting its value for rational biosensor design.

## DISCUSSION

The design of RNA-based biosensors such as toehold switches has long relied on rational engineering and thermodynamic predictions, yet the complexity of RNA folding and context-dependent regulation has limited the accuracy of these approaches. Our study addresses this challenge by combining mechanistic insight with a machine-learning framework to substantially improve predictive performance and design success. We demonstrate that while individual thermodynamic or structural features show only modest correlations with functional outputs, their integration using multivariate modelling by PLS-DA yields a powerful predictive tool capable of robustly classifying high- and low-performing switches across multiple experimental metrics.

Despite other work showing rational features alone are insufficient for accurate toehold switch prediction—based on modest regression performance (r^2^ = 0.20-0.35) and limited classification accuracy (AUROC = 0.76) using multilayer perceptron models (34, 35)—our study demonstrates that these features can, in fact, yield high predictive power when used in a carefully tuned PLS-DA framework. By combining recursive feature elimination, cross-validation, and component optimization, our PLS-DA approach achieved classification accuracies exceeding 90% across multiple performance metrics. While neural networks may suffer from information bottlenecks or overfitting when trained on limited thermodynamic summaries, PLS-DA can effectively extract and leverage multivariate patterns from mechanistic features when the feature space is curated.

By coupling mechanistic insight with a data-driven, interpretable model, we show that rational features remain highly informative when applied with appropriate modelling strategies. Toehold-VISTA is thus an important advance for riboregulator engineering, providing a design tool that is transparent, modular, high-throughput compatible, and grounded in biophysical properties. As RNA diagnostics and synthetic regulatory systems continue to expand, such tools will be essential for bridging the gap between design and function in increasingly complex biological systems.

In addition to advancing predictive modeling for toehold switch performance, our dataset provides a resource for identifying general principles of optimal binding site selection on structured RNAs. Across >200 systematically tiled sites on the mCherry transcript, we observed consistent enrichment of high-performing sites in regions characterized by low proximal base-pairing probability, balanced GC/AU stem composition, and favorable local MFE values when including short flanking regions (±10–25 nt). These features were more predictive of activation than global target folding metrics, underscoring that localized accessibility and moderate stem stability—rather than extreme single-strandedness or rigid base pairing—enable robust strand invasion and hairpin disruption. The mechanistic requirement for an accessible nucleation site followed by a moderately structured downstream segment appears to balance efficient trigger binding with maintenance of a tight OFF state, a trade-off likely relevant to other RNA sensing modalities.

Our findings emphasize that sensor binding site effectiveness is strongly influenced by both proximal and distal features of the target RNA, with proximal structural accessibility dominating the functional response (Figure 3). This insight has broader implications for RNA-targeting technologies beyond toehold switches. For mRNA-binding sensors such as CRISPR-Cas13 effectors (19), ADAR-based RNA editors (5, 6), and miRNA sensors (30, 32, 54, 55), the local RNA structure and sequence context near the binding site likely play a pivotal role in modulating sensor access and efficacy. Proximal secondary structure may occlude guide or sensor binding, while flanking codon usage and translation kinetics may affect local RNA dynamics, influencing the availability of single-stranded regions necessary for efficient interaction. Thus, although our experimental system used toehold switches as the readout, the physical constraints we identify—short-range accessibility, balanced GC content, and avoidance of ribosome-protected regions—likely define generally favorable binding environments on folded RNAs. Incorporating these principles into computational pipelines, as implemented in VISTA, can guide the rational placement of binding sites for new sensors, RNA editors, or riboregulators on any structured transcript of interest. By explicitly linking site-specific structural properties to functional outcomes, our work provides a framework for translating single-target mapping studies into broadly applicable design rules for RNA-targeting technologies.

Moreover, the observed codon usage effects suggest that ribosome-mediated remodeling of RNA structure is a crucial consideration in sensor design. Low codon usage regions may induce ribosomal pausing and transient occlusion of target sites, thereby reducing sensor accessibility (Figure 4). This phenomenon could impact RNA-targeting therapeutics and synthetic biology applications that depend on efficient RNA binding or editing. Previous studies have demonstrated that translation elongation rates modulate RNA folding landscapes and accessibility (37, 38), consistent with our data. Integrating such dynamic cellular features into sensor design frameworks could improve performance across diverse RNA-targeting modalities.

Taken together, these results advocate for a generalizable, target-aware machine learning framework like VISTA that incorporates both sensor and target contextual features to optimize RNA-based sensor design. Future extensions of this approach to ADAR and CRISPR-Cas systems could similarly benefit from mechanistic feature integration and machine learning-guided optimization, enabling broad applicability in RNA synthetic biology and therapeutic development.

## DATA AVAILABILITY

Toehold-VISTA is open source available in the GitHub repository: https://github.com/AlexGreenLab/vista

## SUPPLEMENTARY DATA

Supplementary Data are available at NAR online.

## ACKNOWLEDGEMENTS

This work was supported by startup and DAMP Laboratory Pilot Project funds from Boston University; a National Institutes of Health (NIH) U01 award (1U01AI148319), R01 award (1R01EB031893), and an NIH Director’s Transformative Research Award (R01EB037112) to A.A.G; National Institute of Health Training Program in Synthetic Biology and Biotechnology award (1T32GM130546) and a National Science Foundation Graduate Research Fellowship (2234657) to J.M.R. The content is solely the responsibility of the authors and does not necessarily represent the official views of the National Institutes of Health.

