## Supplementary Information for "Toehold-VISTA: A machine learning approach to decipher programmable RNA sensor-target interactions"

This PDF file includes:

Supplementary Figures 1 to 7

Supplementary Tables 1 to 2

Additional Supplementary Tables:

Supplementary Table 3: Toehold switch and target sequences with experimental measurements

Supplementary Table 4: Calculated biophysical parameters

Supplementary Table 5: Pearson r correlations between biophysical parameters and experimental measurements

Supplementary Table 6: Pairwise probabilities for mCherry RNA

Supplementary Table 7: VISTA-model ranked predictions

Supplementary Table 8: SARS-CoV-2 toehold switch and target sequences with experimental measurements

Supplementary Table 9: Green et al. 2014 mCherry toehold switch benchmark measurements

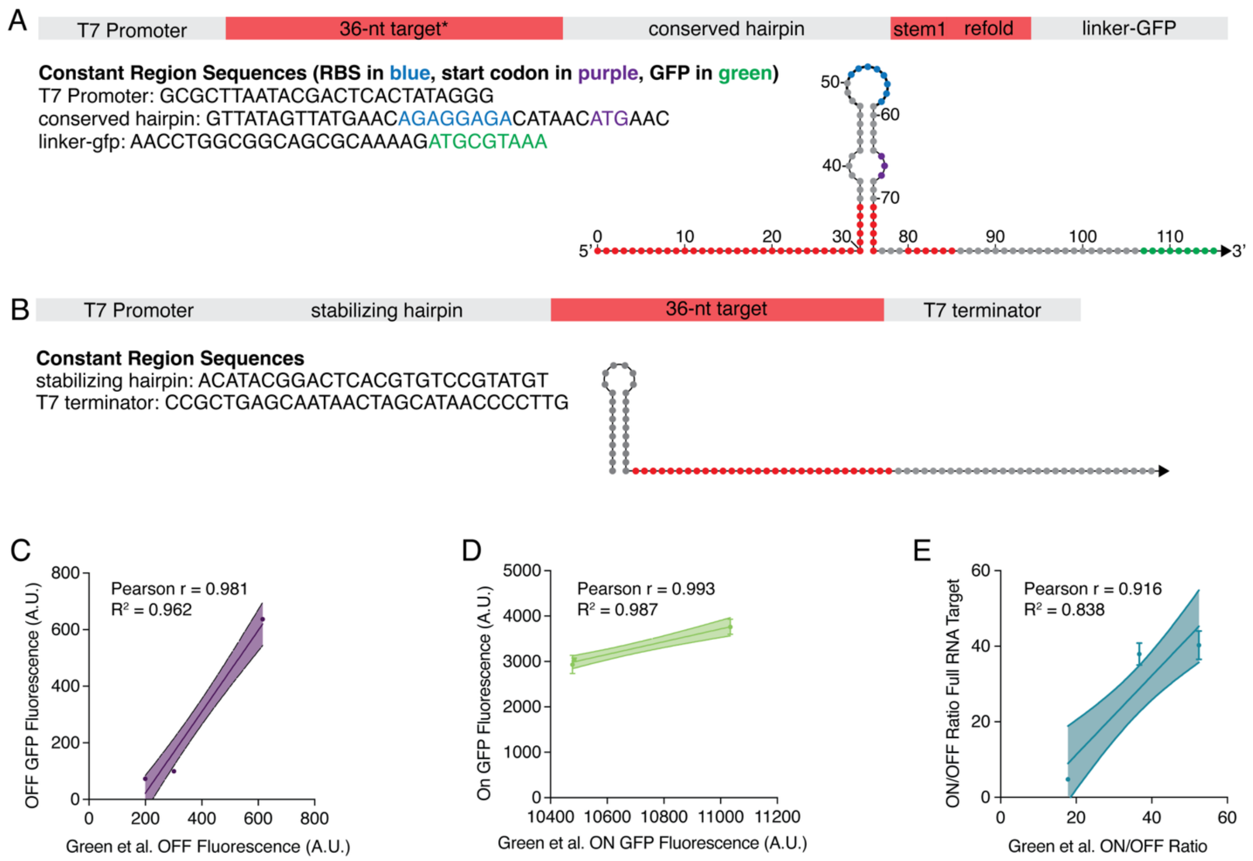

**Supplementary Figure 1. Design and validation of toehold switch library.** Toehold switch constructs were synthesized as single-stranded DNA and representative switches were verified against previously published switches targeting the *mCherry* mRNA. (A) A schematic of a representative toehold switch construct used for the synthesis of each switch RNA. Toehold switch construct regions include the *promoter* (T7 promoter including GGG region), *36-nucleotide target complement* (individual, toehold-unique), *conserved hairpin* (selected from the tsgen2 (1, 2) architecture), *stem1* (complement to the last 6 nucleotides of the trigger binding domain), *refold* (sequesters the bottom part of the toehold switch hairpin after target binding), and *linker-GFP* (21 nucleotide sequence of unstructured amino acids and the first three codons of mut3b-GFP-asv reporter). Further structural details may be found in Supplementary Table 2. (B) Truncated trans RNA target construct design featuring the *promoter* (T7 promoter with GGG region), *stabilizing hairpin* (to stabilize and limit interactions of truncated target with surrounding sequence), *36-nucleotide target* (individual toehold-unique), and *T7 terminator* (first 30 nucleotides of the T7 terminator). The OFF-state GFP fluorescence, ON-state GFP fluorescence, and fold change of three toehold switches targeting mCherry previously characterized by Green et al. (3) were compared against their three closest respective tiles ordered in this dataset: (C) OFF-state GFP expression comparison (Pearson r = 0.981, $r^2 = 0.962$, error bands indicate 95% confidence interval), (D) the ON-state GFP expression comparison (Pearson r = 0.993, $r^2 = 0.987$, error bands indicate 95% confidence interval), and (D) the ON/OFF fold change in GFP expression comparison (Pearson r = 0.916, $r^2 = 0.838$, error bands indicate 95% confidence interval).

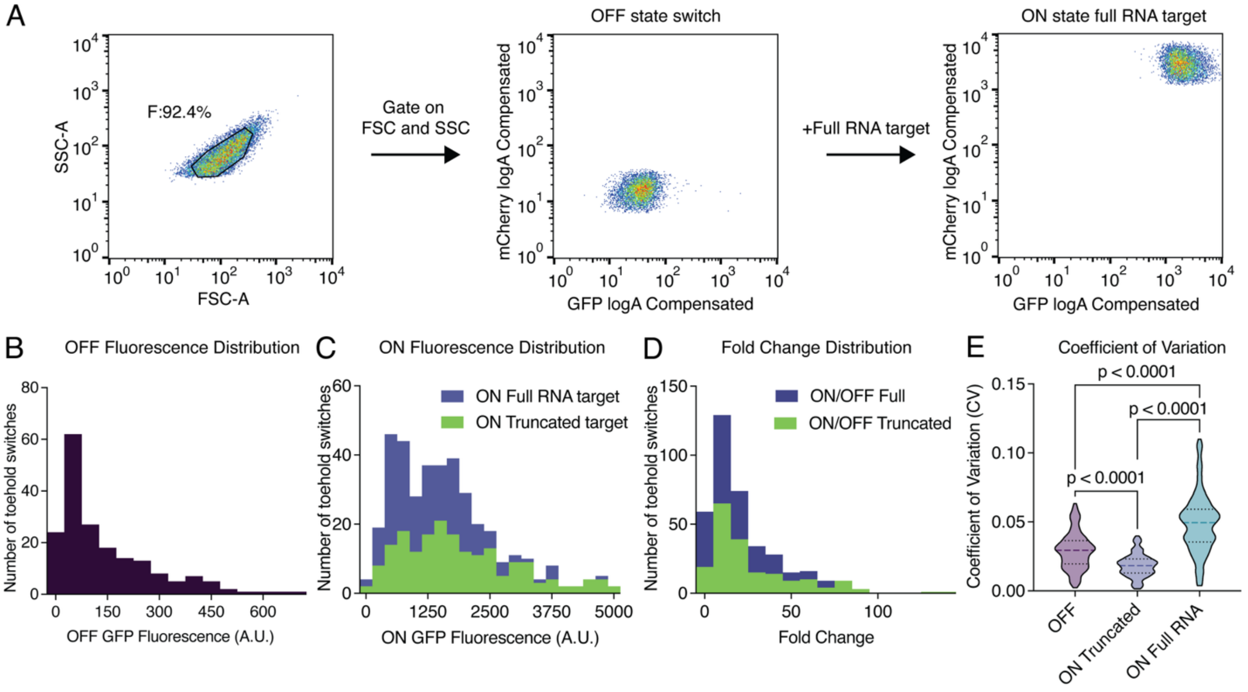

**Supplementary Figure 2. Flow cytometry gating strategy and fluorescence distributions.** (A) The gating strategy for IPTG-induced *E. coli* BL21 (DE3) cells is shown. First, cells were gated by their SSC-A and FSC-A distributions. Populations exhibited a unimodal distribution in GFP-logA and mCherry-logA channels. Measurements of OFF-state GFP fluorescence with non-cognate target RNA (B), as well as ON-state GFP fluorescence in presence of truncated or full-length RNA targets (C) and their respective fold changes (D). Coefficient of variation (CV) for fluorescence characterization of each toehold switch variant (E). CV was determined by dividing the standard deviation and the mean of fluorescence measurements for each toehold switch:target variant, derived from each flow cytometry fluorescence characterization described in (B) and Figure 1B. Comparisons were made by one-way ANOVA. For violin plots, the horizontal dashed line represents the median and dotted lines are at 25$^{th}$ and 75$^{th}$ percentiles.

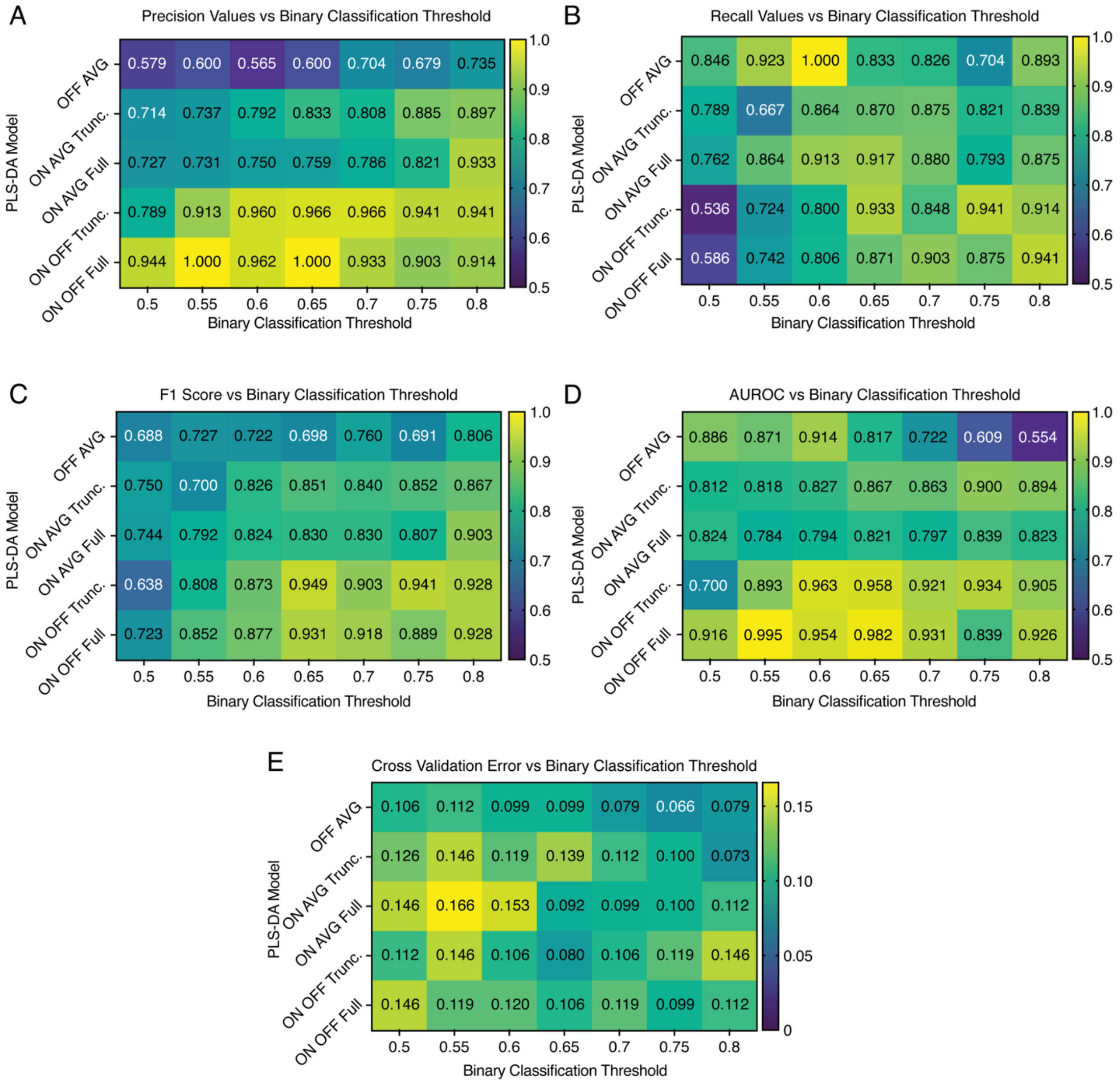

**Supplementary Figure 3. Determination of the optimal ON/OFF binary classification threshold.** Precision (predicted positives that are true positives of "high" class), recall (sensitivity or true positive rate), F1 score (harmonic mean of precision and recall), and area under the receiver operator curve (AUROC) were used to determine the optimal cutoff threshold at which to binarize data for each PLS-DA model. We trained a PLS-DA model on each of the five empirically determined metrics at seven different binarization threshold and compared the following performance metrics: (A) precision results, (B) recall results, (C) F1 score results, and (D) AUROC results. The final threshold selected for the five classification models was 0.65, specifically for balancing a high precision and recall with the best average AUROC for the truncated and full-RNA target fold change models. We also report the cross-validation error (E), a key metric in the validity of a PLS-DA model, for each binary classification threshold and PLS-DA model trained.

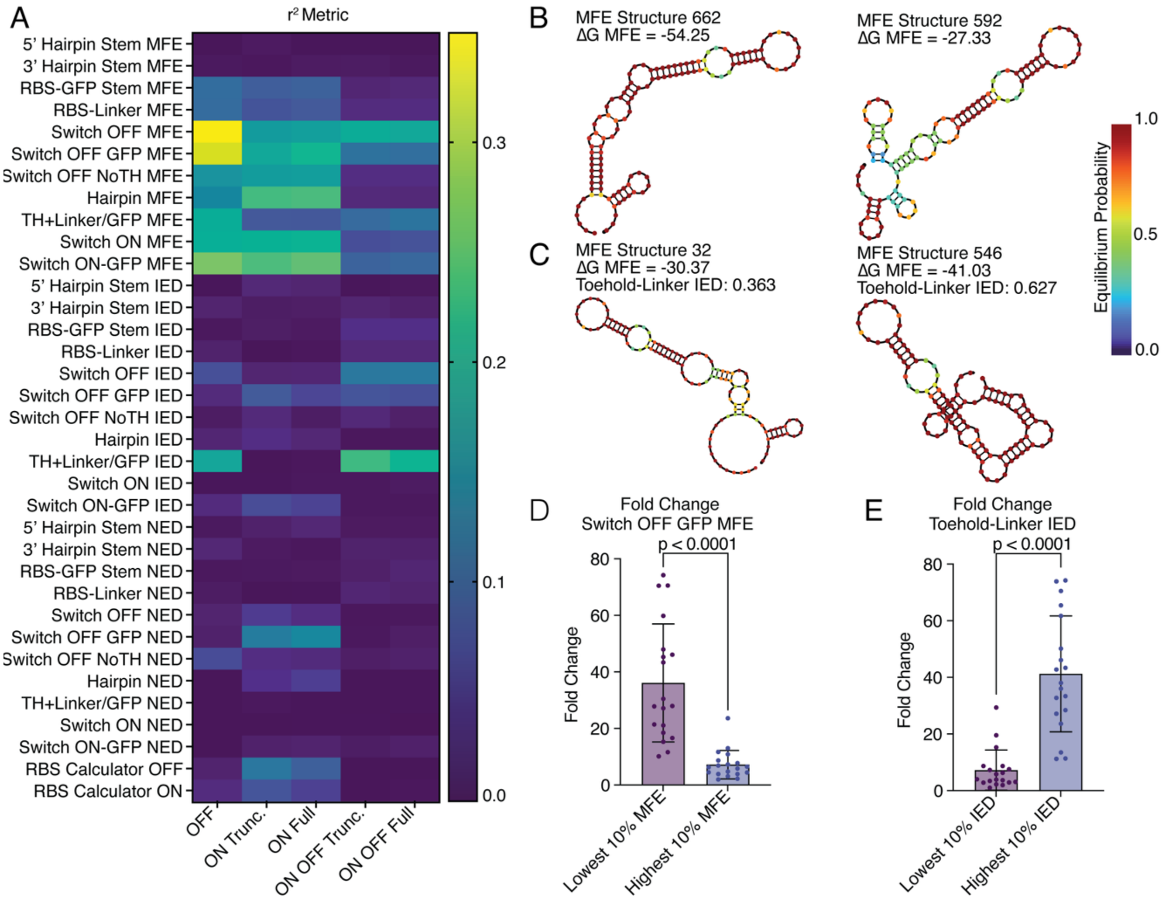

**Supplementary Figure 4. Correlations between rational features and the toehold switch dataset.** (A) Squared Pearson correlation coefficients (max($r^2$) = 0.35) from Figure 2A. (B) Representative NUPACK-determined minimum free energy (MFE) structures for the lowest (switch 662) and highest (switch 592) MFE values for the Switch-OFF MFE metric. (C) Representative NUPACK-determined MFE structures for the lowest (switch 32) and highest (switch 546) toehold-linker/GFP ideal ensemble defect (IED). In each case, the region preceding the hairpin stem should be totally unbound, as shown by the ideal structure in Supplementary Figure 1A. (D) Comparison of the fold change for full RNA target when stratified by the lowest and highest 10% MFE for the "Switch OFF GFP" metric (n =19, $p < 0.0001$ two-tailed t test). (E) Comparison of full RNA target fold change when stratified for lowest and highest 10% IED for the "Toehold linker/GFP" metric. For all bar plots, horizontal line indicates the median and whiskers indicate the standard deviation (SD).

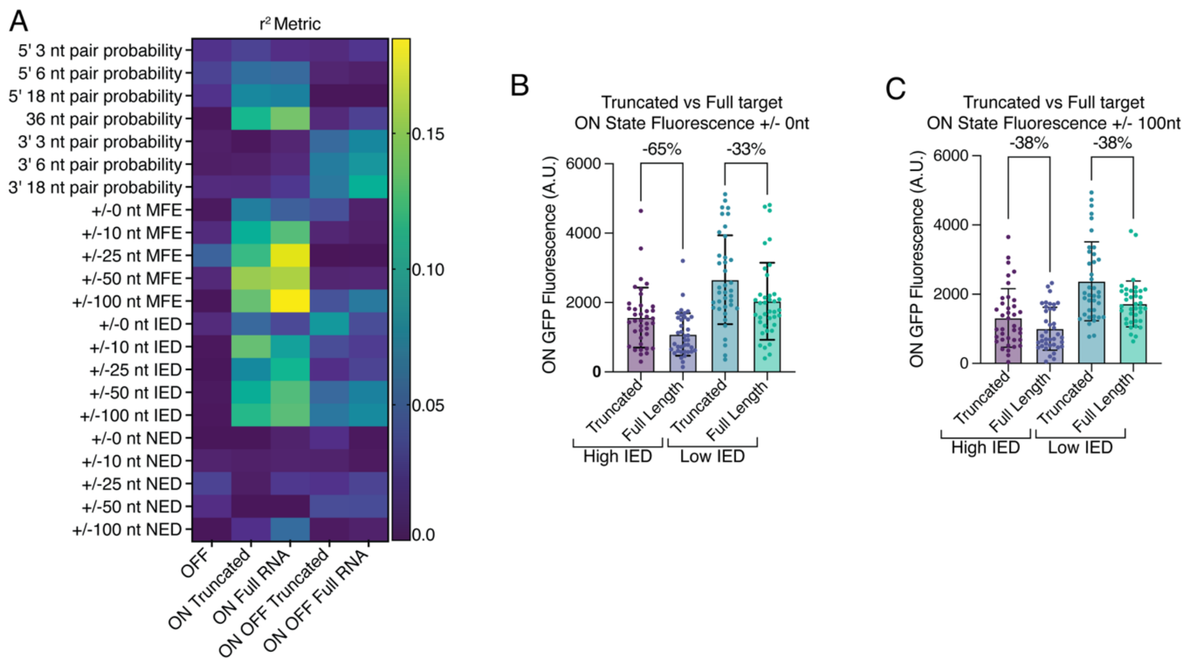

**Supplementary Figure 5. Correlations between rational features of the full RNA target and the toehold switch dataset.** (A) Squared Pearson correlation coefficients (max($r^2$) = 0.18) of correlations from Figure 3B. (B) Truncated and full-length ON-state GFP fluorescence were stratified for the highest and lowest 20% ideal ensemble defect (IED) metric for 0 nucleotide flanking region of the full-length target (n = 38). (C) Truncated and full-length ON-state GFP fluorescence were stratified for the highest and lowest 20% IED metric for 100-nucleotide flanking region of the full-length target (n=38). For all bar plots, horizontal line indicates the median and whiskers indicate the standard deviation (SD).

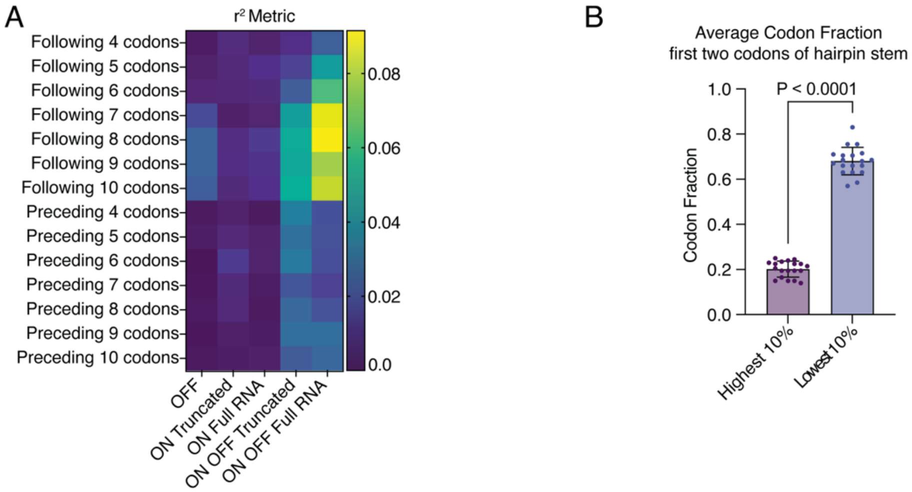

**Supplementary Figure 6. Correlations between codon fraction around full RNA target and toehold switch dataset.** (A) Squared Pearson correlation coefficients (max($r^2$) = 0.09) from Figure 3B. (B) Lowest and highest 10% of average codon fractions in the first two codons at the base of toehold switch hairpin stem (n=19, $p < 0.0001$, two-tailed t test). For all bar plots, horizontal line indicates the median and whiskers indicate the standard deviation (SD).

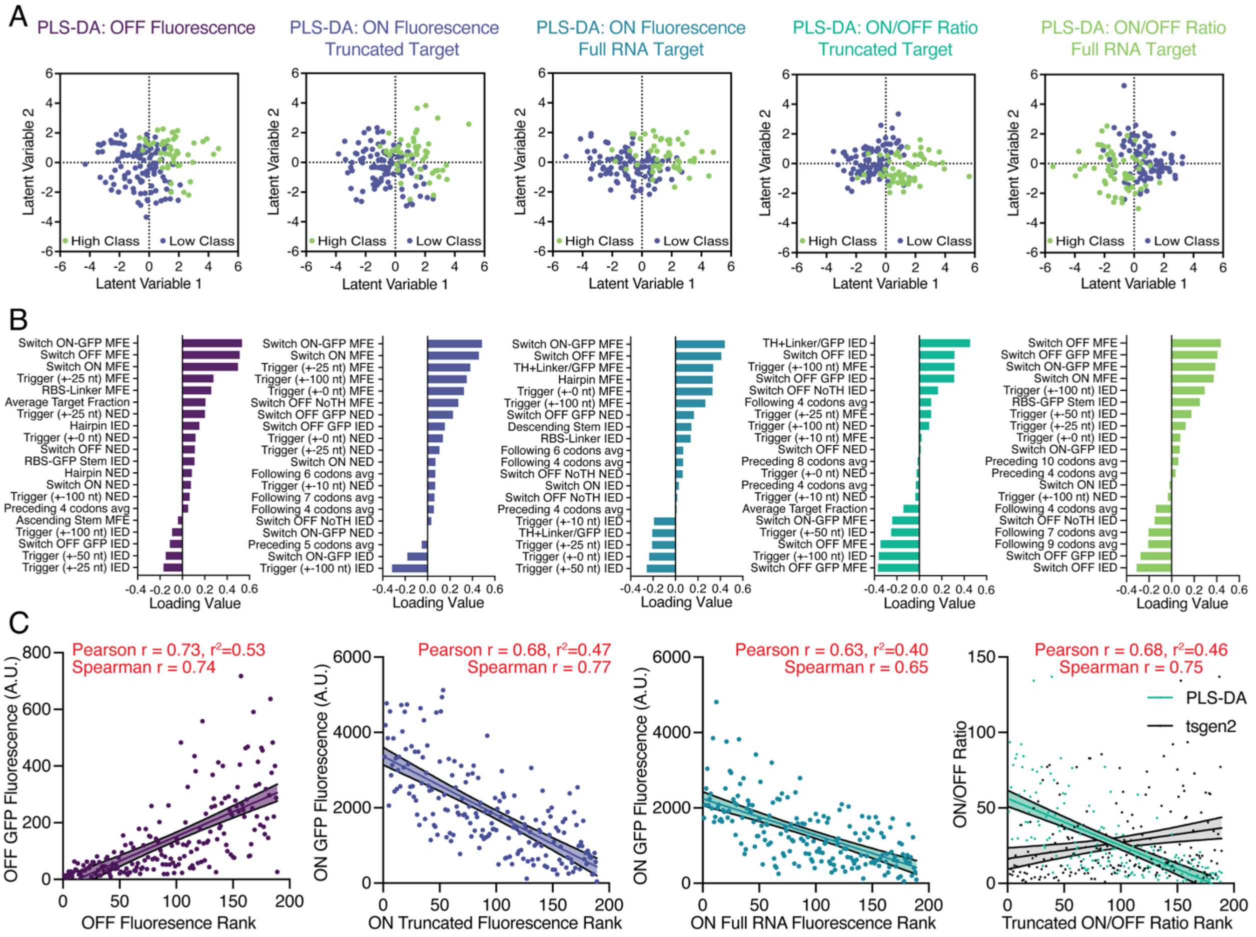

**Supplementary Figure 7. Feature contributions and validation of PLS-DA models.** (A) Latent variable scores plot for each of the five independently trained PLS-DA models. (B) PLS-DA loading plots showing feature contributions to the first latent variable for each performance model. Features with the highest absolute loadings indicate the most influential rational parameter in classifying toehold switch performance. (C) Experimental values plotted against PLS-DA algorithm rank for OFF-state GFP fluorescence, ON-state truncated GFP fluorescence, ON-state full RNA target GFP fluorescence, and truncated ON/OFF fold-change ratio. Linear regression is plotted with 95% confidence interval.

| **Primer Name** | **Sequence** | **Primer Function** |
|---|---|---|
| Switch_BB_fwd | AACCTGGCGGCAGCGCAAAAGATGCGTAAAGGA<br>GAAGAACTTTTCACT | Used to linearize ColA plasmid backbone |
| Switch_BB_rev | TGTTGGGGTTCTCTTAGCTTTGTTTCGCCGCATAA<br>GGGAGAGCGTCGAGATC | Used to linearize ColA plasmid backbone |
| Trigger_BB_fwd | CCGCTGAGCAATAACTAGCATAACC | Used to linearize ColE1 plasmid backbone |
| Trigger_BB_rev | GAGCTATATCGCGAACCACTGGCAGACTACCGA<br>GATCTCGATCCTCTACGC | Used to linearize ColE1 plasmid backbone |
| Switch_insert_fwd | CGGCGAAACAAAGCTAAGAGAACCCCAACAGCG<br>CTAATACGACTCACTATAGGG | Used to amplify toehold switch inserts for ColA backbone |
| Switch_insert_rev | TTTACGCATCTTTTGCGCTGCCGCCAGGTT | Used to amplify toehold switch inserts for ColA backbone |
| Trigger_insert_fwd | GTAGTCTGCCAGTGGTTCGCGATATAGCTCGCG<br>CTAATACGACTCACTATAGGG | Used to amplify 36 nucleotide truncated targets for ColE1 backbone |
| Trigger_insert_rev | CAAGGGGTTATGCTAGTTATTGCTCAGCGG | Used to amplify 36 nucleotide truncated targets for ColE1 backbone |
| GFPmut3b-ASV | ATGCGTAAAGGAGAAGAACTTTTCACTGGAGTTG<br>TCCCAATTCTTGTTGAATTAGATGGTGATGTTAAT<br>GGGCACAAATTTTCTGTCAGTGGAGAGGGTGAAG<br>GTGATGCAACATACGGAAAACTTACCCTTAAATTT<br>ATTTGCACTACTGGAAAACTACCTGTTCCGTGGC<br>CAACACTTGTCACTACTTTCGGTTATGGTGTTCAA<br>TGCTTTGCGAGATACCCAGATCACATGAAACAGC<br>ATGACTTTTTCAAGAGTGCCATGCCCGAAGGTTA<br>CGTACAGGAAAGAACTATATTTTTCAAAGATGACG<br>GGAACTACAAGACACGTGCTGAAGTCAAGTTTGA<br>AGGTGATACCCTTGTTAATAGAATCGAGTTAAAAG<br>GTATTGATTTTAAAGAAGATGGAAACATTCTTGGA<br>CACAAATTGGAATACAACTATAACTCACACAATGT<br>ATACATCATGGCAGACAAACAAAAGAATGGAATC<br>AAAGTTAACTTCAAAATTAGACACAACATTGAAGA<br>TGGAAGCGTTCAACTAGCAGACCATTATCAACAA<br>AATACTCCGATTGGCGATGGCCCTGTCCTTTTAC<br>CAGACAACCATTACCTGTCCACACAATCTGCCCT<br>TTCGAAAGATCCCAACGAAAAGAGAGACCACATG | Used as toehold switch reporter |

| | | |
|---|---|---|
| | GTCCTTCTTGAGTTTGTAACCGCTGCTGGGATTA<br>CACATGGCATGGATGAACTATACAAAAGGCCTGC<br>AGCAAACGACGAAAACTACGCTGCATCAGTTTAA<br>TAA | |
| mCherry | ATGGTGAGCAAGGGCGAGGAGGATAACATGGCC<br>ATCATCAAGGAGTTCATGCGCTTCAAGGTTCACA<br>TGGAGGGCTCCGTGAACGGCCACGAGTTCGAGA<br>TCGAGGGCGAGGGCGAGGGCCGCCCCTACGAG<br>GGCACCCAGACCGCCAAGCTGAAGGTGACCAAG<br>GGTGGCCCCCTGCCCTTCGCCTGGGACATCCTG<br>TCCCCTCAGTTCATGTACGGCTCCAAGGCCTACG<br>TGAAGCACCCCGCCGACATCCCCGACTACTTGA<br>AGCTGTCCTTCCCCGAGGGCTTCAAGTGGGAGC<br>GCGTGATGAACTTCGAGGACGGCGGCGTGGTGA<br>CCGTGACCCAGGACTCCTCCCTGCAAGACGGCG<br>AGTTCATCTACAAGGTGAAGCTGCGCGGCACCA<br>ACTTCCCCTCCGACGGCCCCGTAATGCAGAAGA<br>AGACTATGGGCTGGGAGGCCTCCTCCGAGCGGA<br>TGTACCCCGAGGACGGCGCGCTGAAGGGCGAG<br>ATCAAGCAGAGGCTGAAGCTGAAGGACGGCGGC<br>CACTACGACGCTGAGGTCAAGACCACCTACAAG<br>GCCAAGAAGCCCGTGCAACTGCCCGGCGCGTA<br>CAACGTCAACATCAAGTTGGACATCACCTCCCAC<br>AACGAGGACTACACCATCGTGGAACAGTACGAA<br>CGCGCCGAGGGCCGCCACTCCACCGGCGGCAT<br>GGACGAGCTGTACAAGTAA | Used as full RNA target |
| SARS-CoV-2<br>Nucleocapsid<br>Protein | TATACCATGGGCAGCAGCATGTCTGATAATGGAC<br>CCCAAAATCAGCGAAATGCACCCCGCATTACGTT<br>TGGTGGACCCTCAGATTCAACTGGCAGTAACCAG<br>AATGGAGAACGCAGTGGGGCGCGATCAAAACAA<br>CGTCGGCCCCAAGGTTTACCCAATAATACTGCGT<br>CTTGGTTCACCGCTCTCACTCAACATGGCAAGGA<br>AGACCTTAAATTCCCTCGAGGACAAGGCGTTCCA<br>ATTAACACCAATAGCAGTCCAGATGACCAAATTG<br>GCTACTACCGAAGAGCTACCAGACGAATTCGTG<br>GTGGTGACGGTAAAATGAAAGATCTCAGTCCAAG<br>ATGGTATTTCTACTACCTAGGAACTGGGCCAGAA<br>GCTGGACTTCCCTATGGTGCTAACAAAGACGGCA<br>TCATATGGGTTGCAACTGAGGGAGCCTTGAATAC<br>ACCAAAAGATCACATTGGCACCCGCAATCCTGCT<br>AACAATGCTGCAATCGTGCTACAACTTCCTCAAG<br>GAACAACATTGCCAAAAGGCTTCTACGCAGAAGG<br>GAGCAGAGGCGGCAGTCAAGCCTCTTCTCGTTC<br>CTCATCACGTAGTCGCAACAGTTCAAGAAATTCA<br>ACTCCAGGCAGCAGTAGGGGAACTTCTCCTGCT<br>AGAATGGCTGGCAATGGCGGTGATGCTGCTCTT<br>GCTTTGCTGCTGCTTGACAGATTGAACCAGCTTG<br>AGAGCAAAATGTCTGGTAAAGGCCAACAACAACA<br>AGGCCAAACTGTCACTAAGAAATCTGCTGCTGAG<br>GCTTCTAAGAAGCCTCGGCAAAAACGTACTGCCA<br>CTAAAGCATACAATGTAACACAAGCTTTCGGCAG<br>ACGTGGTCCAGAACAAACCCAAGGAAATTTTGGG | Used in Figure 5 for full-length RNA target |

| | |
|---|---|
| | GACCAGGAACTAATCAGACAAGGAACTGATTACA AACATTGGCCGCAAATTGCACAATTTGCCCCCAG CGCTTCAGCGTTCTTCGGAATGTCGCGCATTGGC ATGGAAGTCACACCTTCGGGAACGTGGTTGACCT ACACAGGTGCCATCAAATTGGATGACAAAGATCC AAATTTCAAAGATCAAGTCATTTTGCTGAATAAGC ATATTGACGCATACAAAACATTCCCACCAACAGA GCCTAAAAAGGACAAAAAGAAGAAGGCTGATGAA ACTCAAGCCTTACCGCAGAGACAGAAGAAACAG CAAACTGTGACTCTTCTTCCTGCTGCAGATTTGG ATGATTTCTCCAAACAATTGCAACAATCCATGAGC AGTGCTGACTCAACTCAGGCCTAA |
| T7 terminator | ccgctgagcaataactagcataaccccttgAGATAACAGATA CttcgGtatctgttatctgttTTTTTTcAACAGATAGCCGCG ttcgCGCGGCtatctgttTTTTTT |

**Supplementary Table 1. Sequences for plasmid construction.** A list of all primers used to construct toehold switch and truncated target plasmids including their name, sequence, and primary function. Also included are reporter protein sequences and the T7 terminator variant used in this study.

| **Rational Feature Name** | **Sequence Region (nucleotide range)** | **Run-length-encoded (RLE) dot-parens-plus and DU+ Notation** | **Description** |
|---|---|---|---|
| 5' hairpin stem | 30:48 | '.18' or<br>U18 | Switch hairpin stem ascending sequence |
| 3' hairpin stem | 58:77 | '.18' or<br>U18 | Switch hairpin stem descending sequence |
| RBS-GFP | 48:116 | '.20 (6 .6 )6 .30' or<br>U20 D6 U6 U30 | RBS sequence to start of GFP |
| RBS-Linker | 48:107 | '.20 (6 .6 )6 .21' or<br>U20 D6 U6 U21 | RBS sequence to end of linker |
| Switch OFF | 0:77 | '.30 (9 .3 (6 .11 )6 .3 )9' or<br>U30 D9 (U3 D6 U11 U3) | OFF conformation of toehold switch |
| Switch OFF-GFP | 0:116 | '.30 (9 .3 (6 .11 )6 .3 )9 .30' or | OFF conformation of toehold switch to GFP start |

| | | | |
|---|---|---|---|
| | | U30 D9 (U3 D6 U11 U3) U30 | |
| Switch OFF No Toehold | 60:116 | ‘(9 .3 (6 .11 )6 .3 )9 .30’ or<br>D9 (U3 D6 U11 U3) U30 | OFF conformation of toehold switch without toehold |
| Hairpin | 30:77 | ‘(9 .3 (6 .11 )6 .3 )9’ or<br>D9 (U3 D6 U11 U3) | Hairpin stem of the toehold switch |
| Toehold + Linker/GFP | 0:30 + 77:116 | ‘.30 + .30’ or<br>U30 + U30 | No hairpin stem; toehold and linker-GFP sequence |
| Switch ON | 0:76 | ‘(36 + )36 .41’ or<br>D36 + U41 | ON conformation of toehold switch to linker |
| Switch ON-GFP | 0:116 | ‘(36 + )36 .6 (6 .11 )6 .3 (6 .6 )6 .30’ or<br>D36 + U6 D6 U11 U3 D6 U6 U30 | ON conformation of toehold switch to GFP |

**Supplementary Table 2. Rational feature sub-sequence regions and description.** Individual sub-regions of the toehold switch from which the 33 rational features were calculated in Figure 2A using NUPACK and ViennaRNA. Sequence nucleotide ranges are reported from the 5’ most position (index 0) to the 3’ most position (index 116); see Supplementary Fig. 1.
